# Gap junctions in the *C. elegans* nervous system regulate ageing and lifespan

**DOI:** 10.1101/657817

**Authors:** Nathalie Alexandra Vladis, Katharina Elisabeth Fischer, Eva Digalaki, Daniel-Cosmin Marcu, Modestos Nakos Bimpos, Peta Greer, Alice Ayres, Qiaochu Li, Karl Emanuel Busch

**Author notes:** these authors contributed equally to this work (Nathalie Alexandra Vladis, Katharina Elisabeth Fischer). Correspondence should be addressed to K.E.B.

## Abstract

The nervous system is a central regulator of longevity, but how neuronal communication interfaces with ageing pathways is not well understood. Gap junctions are key conduits that allow voltage and metabolic signal transmission across cellular networks, yet it has remained unexplored whether they play a role in regulating ageing and longevity. We show that the innexin genes encoding gap junction subunits in *Caenorhabditis elegans* have extensive and diverse impacts on lifespan. Loss of the neural innexin *unc-9* increases longevity by a third and also strongly benefits healthspan. *Unc-9* acts specifically in a glutamatergic circuit linked to mechanosensation. Absence of *unc-9* depends on a functional touch-sensing machinery to regulate lifespan and alters the age-dependent decline of mechanosensory neurons. The life extension produced by removal of *unc-9* requires reactive oxygen species. Our work reveals for the first time that gap junctions are important regulators of ageing and lifespan.

## Introduction

The nervous system is a central regulator of ageing and lifespan^1^. Nerve cells need to sustain their internal homeostasis throughout life to maintain neural function and behavior. However, as their electrical activity is energetically and metabolically demanding, they become vulnerable to the resulting physiological stress^2^. Loss of cellular homeostasis, such as of Ca^2+^ levels, redox balance or energy state, contribute to neuronal dysfunction and neurodegeneration and play an essential role in ageing processes^3^. However, it is not well understood how cellular homeostasis processes interface with and signal to other cells and tissues to regulate ageing in a non-cell-autonomous way. The nexus of how age-dependent cellular and physiological changes impinge on and relate to the processes controlling lifespan also remains largely unexplained^4^.

Gap junctions are unique and ubiquitous intercellular conduits that play fundamental roles in the development and physiology of animals^5, 6^. They are formed by channels on the plasma membrane of two adjacent cells that dock onto each other, allowing the direct transmission of voltage signals between the coupled cells^7^. They functionally join the cytoplasm of the two cells and thus also other small molecules such as second messengers or metabolites can passage through them, allowing biochemical or metabolic coupling^8^. Gap junctions are abundant in the nervous systems where they function as electrical synapses, and are critical for higher-order neural functions such as synchronicity, oscillatory activity or coincidence detection^9^. Conversely, gap junction coupling can also have pathogenic consequences and is responsible for the propagation of cellular injury or death signals to “bystander” cells^10–12^. For example, an injury-mediated increase in neuronal gap junction coupling is part of the mechanism causing excitotoxicity and neuronal death^13^. However, it is currently unknown whether gap junction coupling as a key intercellular communication channel contributes to the regulation of ageing and longevity.

Here we set out to explore if gap junctions affect ageing using the nematode *Caenorhabditis elegans*, an outstanding model for studies on ageing^14, 15^. We assayed the lifespan of loss-of-function mutants of most of the 25 *C. elegans* innexins and discovered that innexins have a significant impact on lifespan, some leading to an extension of lifespan while others shorten it. Surprisingly, most null mutations of neural innexins extended lifespan, with loss of *unc-9*, the most widely expressed innexin in the nervous system, increasing longevity by a third. Its selective removal from glutamatergic neurons led to an increase in lifespan and points to a mechanosensory circuit where UNC-9 regulates ageing. Our results show that UNC-9 alters the age-dependent decline of touch-sensing neurons and depends on functional touch sensation as well as reactive oxygen species to modulate lifespan. This study gives strong evidence to suggest an important and previously unknown role for gap junction intercellular communication in shaping ageing and longevity.

## Results

### The *C. elegans* gap junction genes regulate longevity

As there is extensive intercellular gap junction coupling in all organs of *C. elegans,* we hypothesised that gap junction channels may play specific roles in organismal ageing and longevity. The channel subunits encoded by 25 innexin genes show diverse, highly combinatorial, plastic and dynamic expression in virtually all organs and cells of this animal. The gap junction channels these innexins form play key roles in intercellular communication^16–18^. We asked if innexins influence longevity, which has not been known. To this end we performed lifespan assays in loss-of-function mutants of all innexins except *inx-3*, *inx-12* and *inx-13*; these genes are essential for embryonic development or osmoregulation and their loss confers a lethal phenotype^18, 19^. Loss-of-function mutants are available for all remaining 22 innexins and most of them are putative null alleles (Supplementary Table 1). Lifespan assays were conducted under standard conditions at 20°C. We found that innexins have profound and distinct impacts on lifespan (Fig. 1; Fig. 2; Supplementary Table 2; Supplementary Fig. 1). The effect of the mutants on lifespan differed widely, ranging from *inx-6*, which reduced lifespan by a third, to *unc-9*, which expanded it by the same length. Not only the magnitude of the effects differed, but also their shape; for example, loss of *inx-2* generally shifted the survival curve to the right, while the positive effect of the *inx-15* mutation on lifespan was seen predominantly in the long-lived segment of the population (Fig. 1e,o). Four innexin mutants, *inx-5*, *inx-7*, *inx-10* and *inx-16,* had no significant effect on lifespan (Fig. 1; Fig. 2). Six innexin mutants showed reduced lifespan, namely *eat-5*, *che-7*, *inx-6*, *inx-8*, *inx-21* and *inx-22*. This indicates that the functions conferred by these innexins is necessary to achieve normal lifespan in N2 wild-type animals. To our surprise, twelve innexin mutants extended lifespan, constituting the largest group in the assay (Fig. 2). This indicates that the presence of these innexins in *C. elegans* reduces longevity.

**Fig. 1.**
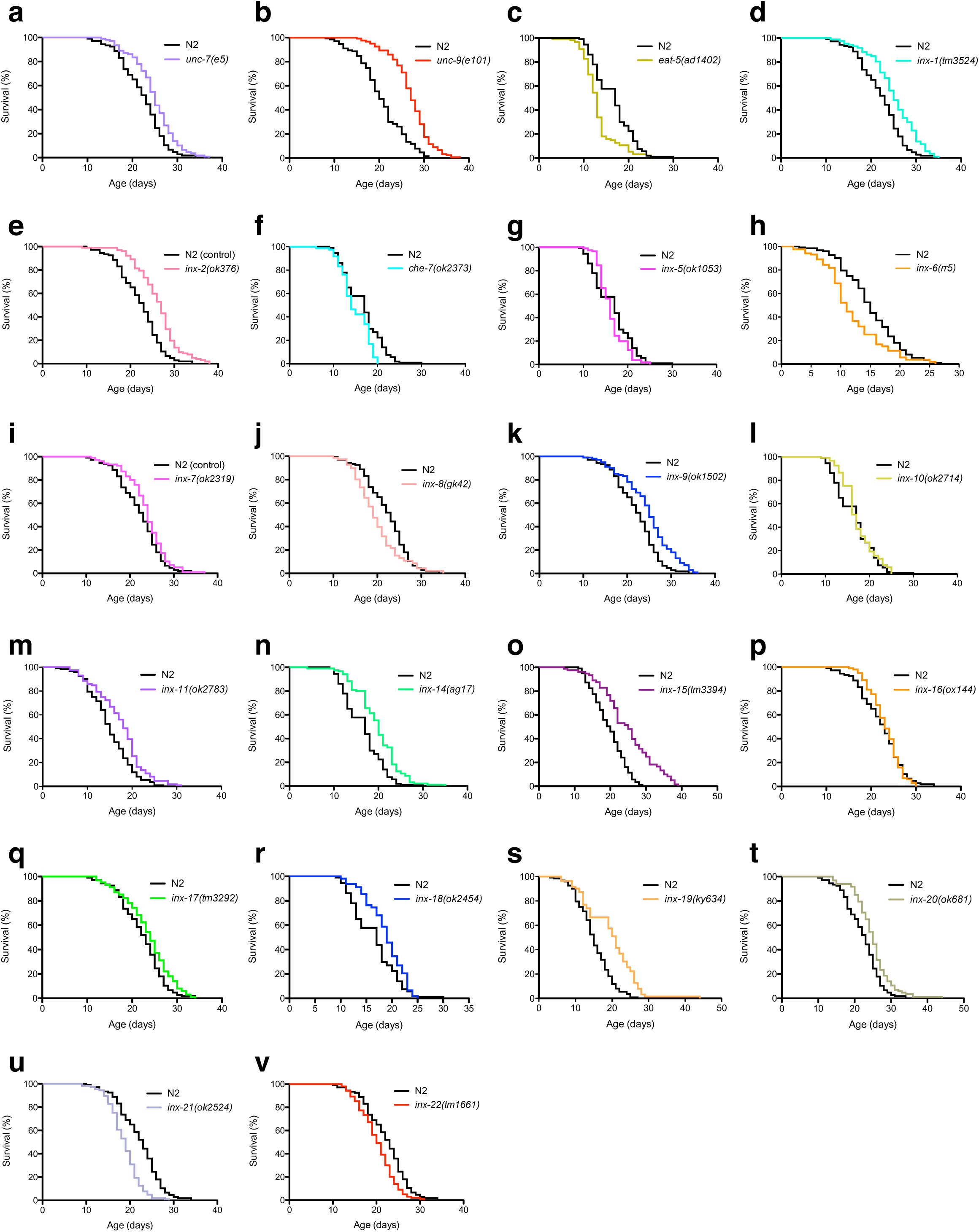
The *C. elegans* gap junction genes regulate longevity. Numbers in non-italicised brackets show number of animals per group for the respective experiment. **a** Lifespan of N2 worms and *unc-7(e5)* (120), (**b**) *unc-9(e101)* (150), (**c**) *eat-5(ad1402)* (130), (**d**) *inx-1(tm3524)* (120), (**e**) *inx-2(ok376)* (120 for N2, 121 for *inx-2*), (**f**) *che-7(ok2373)* (130), (**g**) *inx-5(ok1053)* (130 for N2, 128 for *inx-5*), (**h**) *inx-6(rr5)* (130), (**i**) *inx-7(ok2319)* (120), (**j**) *inx-8(gk42)* (120), (**k**) *inx-9(ok1502)* (120), (**l**) *inx-10(ok2714)* (130), (**m**) *inx-11(ok2783)* (130), (**n**) *inx-14(ag17)* (130), (**o**) *inx-15(tm3394)* (120), (**p**) *inx-16(ox144)* (120), (**q**) *inx-17(tm3292)* (120), (**r**) *inx-18(ok2454)* (130), (**s**) *inx-19(ky634)* (130 for N2, 131 for *inx-19*), (**t**) *inx-20(ok681)* (120), (**u**) *inx-21(ok2524)* (120 for N2, 130 for *inx-20*), (**v**) *inx-22(tm1661)* (120 for N2, 130 for *inx-22*) mutants. For p values see Fig. 2.

**Fig. 2.**
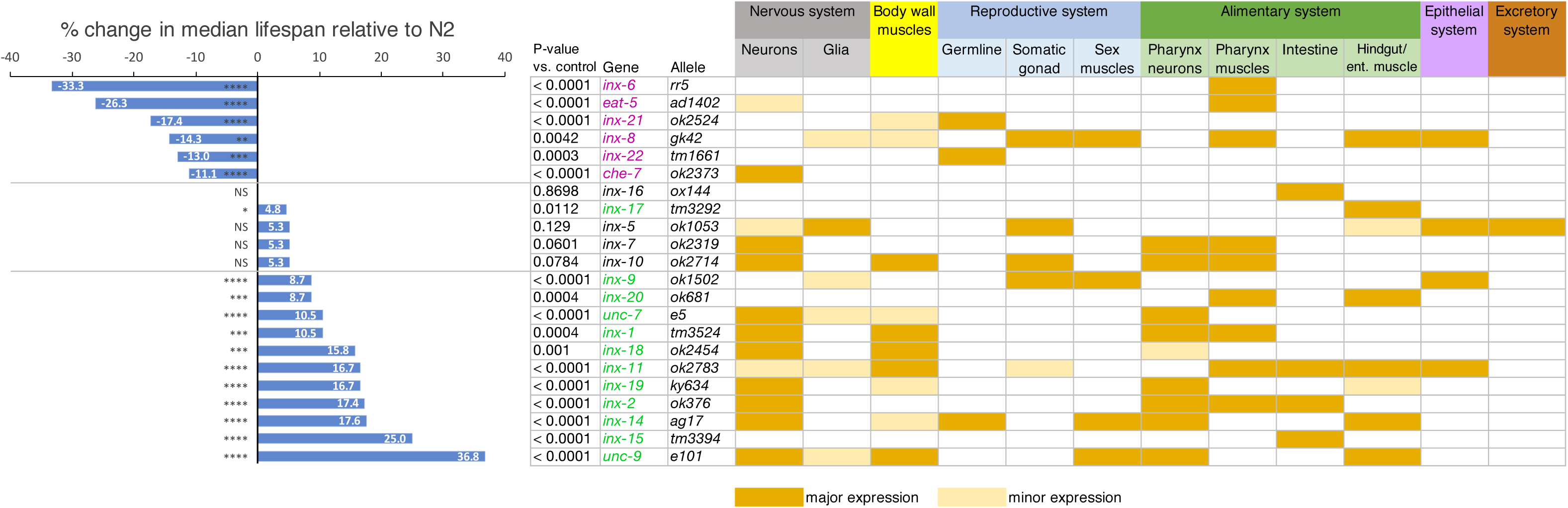
The effects of innexins on lifespan correlate with their tissue expression patterns. Percentage change in lifespan (left) relative to N2 control animals produced by innexin mutations along with normal expression patterns in major *C. elegans* tissues (right). *P<0.05; ***p<0.001; ****p<0.0001; NS not significant, using Log-rank test.

Overall, our results demonstrate that the innexin gene family in *C. elegans* has extensive and diverse effects on lifespan.

### The effects of innexins on lifespan correlate with their tissue expression patterns

Given that the mutants of different innexins have distinct effects on longevity, we sought to compare these effects with the expression patterns and known functions of these genes, based on the literature (Fig. 2). *C. elegans* gap junctions have a plethora of functions in all organs, with different groups of innexins being expressed in each of them^18, 20^.

Some of the most striking effects of innexin mutants were seen in those important for food intake in the pharynx. EAT-5 and INX-6 regulate synchronised muscle contractions in the pharynx and their mutations are defective in feeding ^21, 22^. We found that their mutants have the shortest lifespan observed in the assay (Fig. 2). The *eat-5* mutants developed slowly and remained frail and small until their death. Similarly, *inx-6* mutants had overall poor health.

Apart from the pharyngeal pumping-defective *eat-5* and *inx-6*, mutants of innexins with expression in the alimentary canal generally had increased longevity. The absence of *inx-15*, which is expressed only in the intestine, strongly increased lifespan; no phenotypes had been reported for *inx-15* previously. *Inx-11* and *inx-20* are expressed in the pharynx muscles and hindgut and *inx-11* additionally in the intestine; their mutations also extended lifespan. No defects in pharynx function have been reported for them. INX-16 promotes intestinal muscle contractions by propagating Ca^2+^ waves between cells, and its mutant has defective defecation and constipation^23^; we did not observe a change of lifespan in *inx-16,* however.

The innexins that function in the reproductive system show opposing effects on lifespan. While mutants of two innexins primarily expressed in the germline, *inx-21* and *inx-22*, shortened lifespan, loss of *inx-14* significantly increased it. *Inx-8* and *inx-9*, which are expressed in the somatic gonad, also had opposing effects on lifespan, with the *inx-8* mutant reducing lifespan, while *inx-9* mutants lived longer.

Most remarkably, 8 out of 12 innexin mutants with a positive effect on lifespan are expressed in the nervous system, with intestinal *inx-15* as the primary exception. Conversely, of the innexins with significant neural expression, only *che-7* mutants had a reduced lifespan (Fig. 2). *Che-7* mutants are defective in chemotaxis^24^. *Unc-9* and *unc-7* have several functional roles in the nervous system, including in regulating locomotion, and *inx-19* is required for cell fate determination ^16, 17, 25–28^. All three mutants significantly increased longevity.

The neural innexins show a high degree of overlap with those expressed in the body wall muscles and the pharyngeal nervous system. Of the six innexins that contribute to the electrical coupling of body wall muscles^29^, four mutants show lifespan expansion, namely *inx-1*, *inx-11*, *inx-18* and *unc-9*. All of them also have neural expression. Likewise, the innexins expressed in pharynx neurons are all present in the somatic nervous system as well^20^. This precludes assigning specific roles for innexins in either organ in regulating lifespan. The only innexin widely expressed in glia, *inx-5*, did not affect lifespan. Together, we conclude that despite the highly combinatorial expression of the gap junction gene family in *C. elegans*, the change in lifespan exerted by many innexin mutants aligns with their expression patterns and functional domains.

### *unc-9* regulates longevity in the same pathway as *unc-7* innexin and *unc-1* stomatin

It is highly surprising that loss-function mutants of most innexins expressed in the nervous system prolonged lifespan, as it suggests that the intercellular coupling provided by the gap junctions they form has a negative impact on longevity. To explore this further, we focussed on the innexin *unc-9*, whose canonical putative null mutation *e101*^30, 31^ showed the strongest increase in lifespan in our assay (Fig.1; Fig. 2; Supplementary Table 2). *Unc-9* is expressed in 97 of 104 somatic neuron classes. In addition, it is expressed in pharyngeal neurons and most muscles^18, 20^. To confirm that the long life of *e101* mutants is caused by the absence of *unc-9*, we tested a second allele, *fc16*, a putative null mutation^31^ (Supplementary Table 1). This mutant showed significantly prolonged lifespan as well (Fig. 3a). We also created a transgenic rescue line that bears a fosmid containing the *unc-9* locus in the *e101* mutant background. We confirmed that this line indeed rescues *unc-9* function by placing worms in liquid medium and counting the number of lateral swimming movements (Supplementary Fig. 2). Lifespan of this strain was significantly shorter than that of the mutant and resembled that of the N2 wild-type control, indicating that it is indeed the loss of *unc-9* which increases longevity (Fig. 3b).

**Fig. 3.**
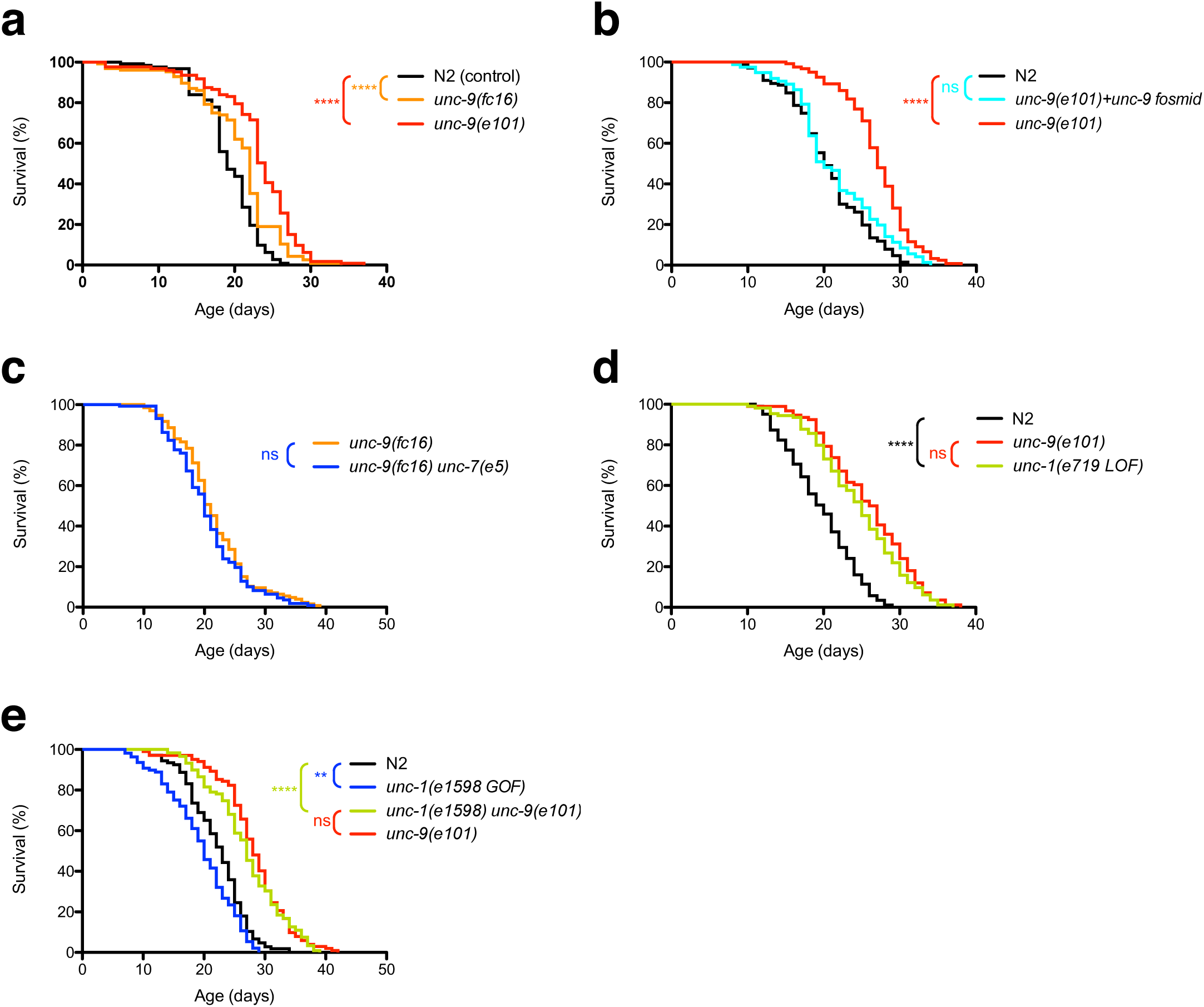
*unc-9* regulates longevity in the same pathway as *unc-7* innexin and *unc-1* stomatin. **a** *unc-9(fc16)* and *unc-9(e101)* mutants increase lifespan relative to N2 controls (N=130 animals per group). **b** Fosmid rescue of *unc-9(e101)* reduces lifespan to N2 control levels (N=150 animals per group). **c** *unc-9(fc16) unc-7(e5)* double mutants have the same lifespan as *unc-9(fc16)* mutants (N=150 animals per group). **d** *unc-1(e719 LOF)* mutants have similarly extended lifespan as *unc-9(e101)* mutants (N=120 animals per group). **e** *unc-9(e1598 GOF)* mutants (N=120 animals) have significantly reduced lifespan compared to N2 controls (N=120 animals). *unc-1(e1598) unc-9(e101)* mutants (N=130 animals) have similar lifespans as *unc-9(e101)* mutants (N=120 animals). **P<0.01; ****p<0.0001; ns not significant, using Log-rank test.

The functions of UNC-9 in the nervous system appear to be closely linked to those of another innexin, UNC-7. UNC-7 has the highest amino acid sequence similarity to UNC-9, and *unc-7* mutants phenocopy the locomotion defect of *unc-9* mutants; both co-localise extensively in the nervous system. UNC-9 and UNC-7 are thought to form heterotypic gap junctions which can act as rectifying electrical synapses, especially in interneuron-motoneuron pairs to generate the undulatory wave in locomotion, and control direction of movement ^25, 26, 32, 33^. We therefore wanted to test if *unc-9* and *unc-7* cooperate in the regulation of longevity and compared the lifespan of *unc-9* single and *unc-9 unc-7* double mutants. As they did not show additive lifespan extension, we suggest that both innexins act in the same pathway to modulate longevity (Fig. 3c).

The stomatin-like integral membrane protein UNC-1 is specifically required for the function of gap junctions formed by UNC-9^34^. *Unc-1* loss of function mutants display the same uncoordinated locomotion phenotype as animals lacking *unc-9*^35^. The protein colocalises with UNC-9 and is thought to regulate the gating of the gap junction channels it forms, with the channels being in a closed state when UNC-1 is absent. We therefore sought to test if the absence of *unc-1* also affects ageing. We found that the putative null mutation *unc-1(e719)* strongly extends lifespan, in a nearly identical manner as the *unc-9* null mutation does (Fig. 3d). In contrast, the dominant gain-of-function *unc-1(e1598)* allele had the opposite effect and significantly reduced the mutant’s longevity compared to N2 (Fig. 3e). If the gain-of-function form of UNC-1 shortens lifespan by acting via UNC-9, we would expect this effect to disappear in a double mutant bearing both the *unc-1* gain-of-function and the *unc-*9 loss-of-function alleles. We observed that lifespan of this double mutant resembles that of *unc-9* mutants alone and is significantly longer than that of both N2 and the *unc-1* gain-of-function mutants (Fig. 3e). Therefore, *unc-9(e101)* is epistatic to *unc-1(e1598)* in the control of lifespan.

These results together support a model where functional intercellular coupling by UNC-9/ UNC-7 gap junction channels reduces the lifespan of *C. elegans*.

### Lack of the unc*-9* innexin also improves healthy ageing of *C. elegans*

Although lifespan is an excellent proxy for the rate of ageing, ageing is a broader process and encompasses also the progressive decline of the physiological functions of an organism over its lifetime^36, 37^. This decline is conveyed by the concept of ‘healthspan’, which is the proportion of life spent in a healthy state^38^. Genetic pathways that modulate lifespan of *C. elegans* have divergent effects on healthspan, with some mutants such as *daf-2* increasing healthspan, while others reduce healthy ageing^39, 40^. We therefore sought to measure whether the mutants of *unc-9* are not only long-lived but also associated with increased healthspan. Because of the predominantly neural expression of *unc-9*, we measured indicators for the progressive functional decline of the nervous system with age, namely locomotion, responsiveness to touch, pharyngeal pumping and defecation.

First, we measured the change in locomotory speed with age, a well-documented indicator for the decline of physical function^38, 41, 42^. To separate the ageing effect of *unc-9* from the general uncoordinated locomotory defect caused by these mutants, we normalised speed relative the maximum locomotory speed observed in day 3 adults, as done previously for *daf-2* mutants, which also affect locomotion^42^. We found that the decline of locomotory speed in ageing worms is significantly delayed in *unc-9* mutants and that the animals perform consistently better than wild type until the end of their lives (Fig. 4a). We used locomotory activity to compare the healthspan (defined as the period with >50% of maximal activity) ratio of *unc-9(e101)* and N2 wild-type worms, following established procedure^40, 43^. Healthspan was extended by 6.4 days in the innexin mutant (11.7d in *unc-9(e101)*, 5.3d in N2), a ratio of 2.2 (Fig. 4a). Our results suggest that loss of *unc-9* increased healthspan more than twofold compared with wild type. As a second comparison, we integrated the areas under the relative locomotory speed curves as an indicator of overall behavioural performance, as done previously^40^; here, *unc-9* mutants showed a 1.8-fold increase over N2 wild-type animals (10.92 vs. 6.19). Thus, according to either measure, the increased relative locomotory speed is considerably higher than the increase in median lifespan (37%).

**Fig. 4.**
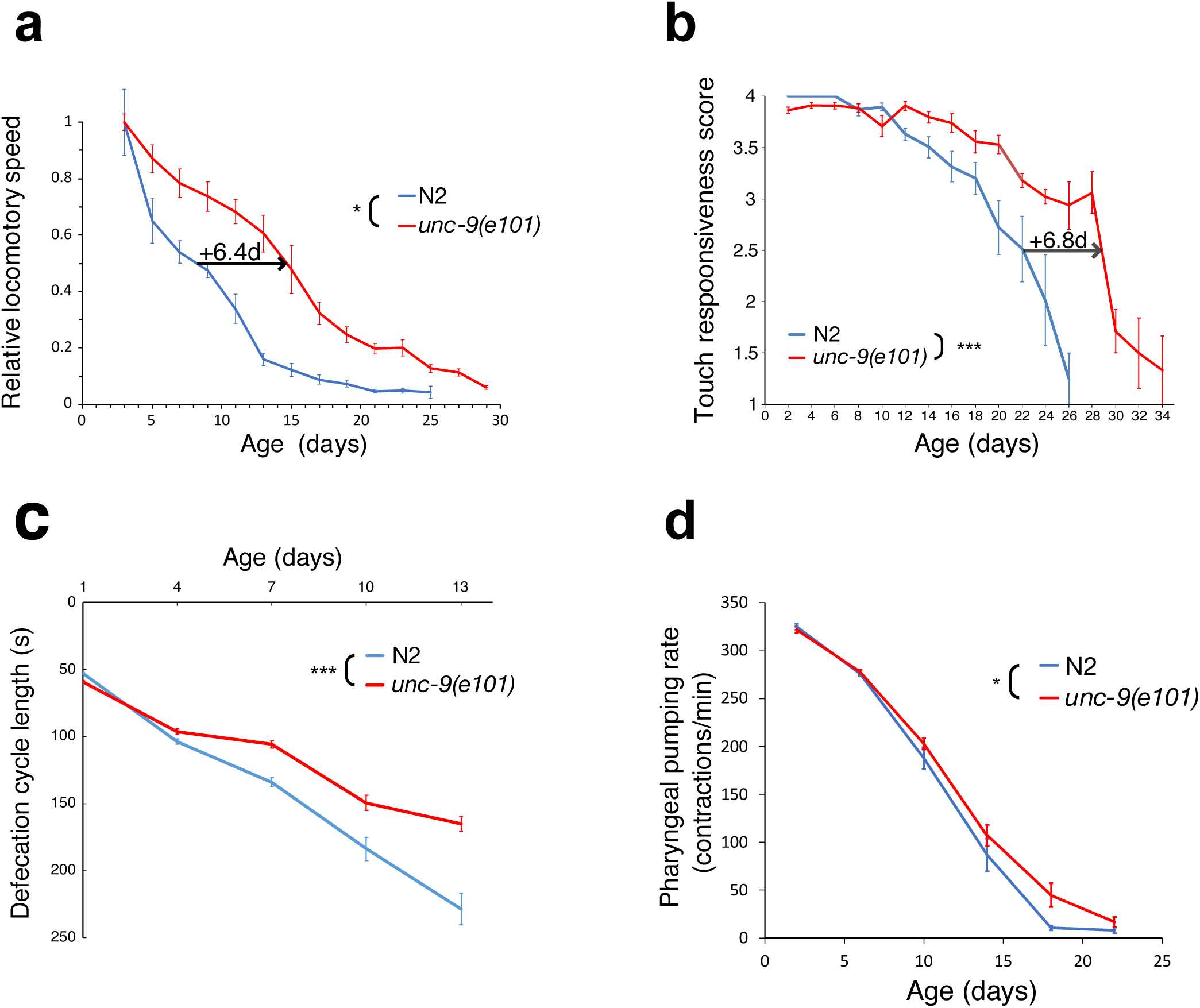
Lacking the *unc-9* innexin also improves healthy ageing of *C. elegans*. **a** Relative locomotory speed declines more slowly with age in *unc-9(e101)* mutants than in N2 controls (N=9 plates per group). The arrow indicates the delay of *unc-9* mutant worms in reaching 50% of maximal functional capacity, which defines healthspan, relative to N2 wild type. **b** Touch responsiveness declines more slowly in ageing *unc-9(e101)* mutants than in N2 (N=120 animals per group). The arrow indicates the delay of *unc-9* mutant worms in reaching 50% of maximal functional capacity relative to N2. **c** Change in defecation cycle duration with age in *unc-9(e101)* mutants and N2 controls (N=26 animals per group). **d** Change in pharyngeal pumping rate in ageing *unc-9(e101)* mutants and N2 controls (N=15 animals per group). *P<0.05; ***p<0.001; ns not significant; (**a**), (**c**), (**d**) mixed effects model; (**b**) generalised estimating equations.

To gauge the qualitative decline of sensory responses with age, we then measured mechanosensory responsiveness of animals to harsh touch. Both N2 wild-type and *unc-9* mutant animals remained fully responsive until day 10. In older worms, the ability to respond to touch was greater and more persistent in the absence of *unc-9*, while N2 declined significantly faster than the innexin mutants (Fig. 4b). Half-maximal responses in the mutant were reached 6.8 days later than in controls, expanding healthspan of this parameter for the same length as the increase in median lifespan (26 vs. 19 days in *unc-9(e101*) and N2).

Another key physiological process that shows age-related dysfunction is defecation. The frequency of defecation cycles, which are highly stereotypic and periodic in young *C. elegans*, reduces significantly in ageing worms^44^. We found that the decline in defecation cycle length is considerably and significantly slower in *unc-9* mutants (Fig. 4c). In young adults, *unc-9* defecate at slightly but significantly lower frequency than N2 (59s vs. 53s, p<0.0001), but at all later time points N2 defecate less frequently.

We also measured pharyngeal pumping, which indicates feeding activity and is also subject to age-dependent decline^45^. In young and middle-aged adults, pumping rates were virtually identical between N2 and *unc-9* mutants (p=0.711 for day 2 adults, n.s., using a linear mixed model with repeated measurements). *unc-9* animals therefore do not have a general defect in bacterial feeding. In old animals, pumping rates declined significantly less in *unc-9*, an effect that increased over time. (Fig. 4d).

We additionally measured changes in the locomotory responses to a tap stimulus with age in *unc-9* and N2 animals but did not observe consistent differences between the mutant and wild type (Supplementary Fig. 3). Since *unc-9* is expressed in the sex muscles and has an egg retention phenotype^18, 31^, we examined if *unc-9* mutants are defective in egg-laying. We found that the brood size of *unc-9(e101)* mutants is 258±7 per parent (n=18), which is 13% lower compared to N2, which has 295±6 offspring per hermaphrodite (n=20). This slightly reduced fertility is unlikely to affect lifespan.

Taken together, these results suggest that electrical coupling by UNC-9 regulates not only lifespan but also the decline of specific outputs of the nervous system with age, and that ageing worms lacking this innexin are more active and responsive than wild type, showing improved healthspan.

### *unc-9* acts in the glutamatergic nervous system to regulate lifespan

The clear effect of UNC-9 on lifespan and healthspan raises the question of where it acts to control longevity. *unc-9* is widely expressed in the nervous system and muscles. To determine if *unc-9* acts in muscles or the nervous system to regulate lifespan, we knocked down *unc-9* in N2 wild-type animals and the RNAi-hypersensitive strain KP3948, using RNA interference by bacterial feeding. In N2 but not KP3948, the nervous system is refractory to RNAi. To avoid developmental defects of *unc-9* knockdown, we started the RNAi treatment at the L4 larval stage. We found that *unc-9* knockdown extends lifespan in KP3948 but not N2 animals, suggesting that the nervous system is the organ where UNC-9 affects lifespan (Fig. 5a,b). This result further corroborates our previous finding using *unc-9* mutants that the absence of *unc-9* causes lifespan extension.

**Fig. 5.**
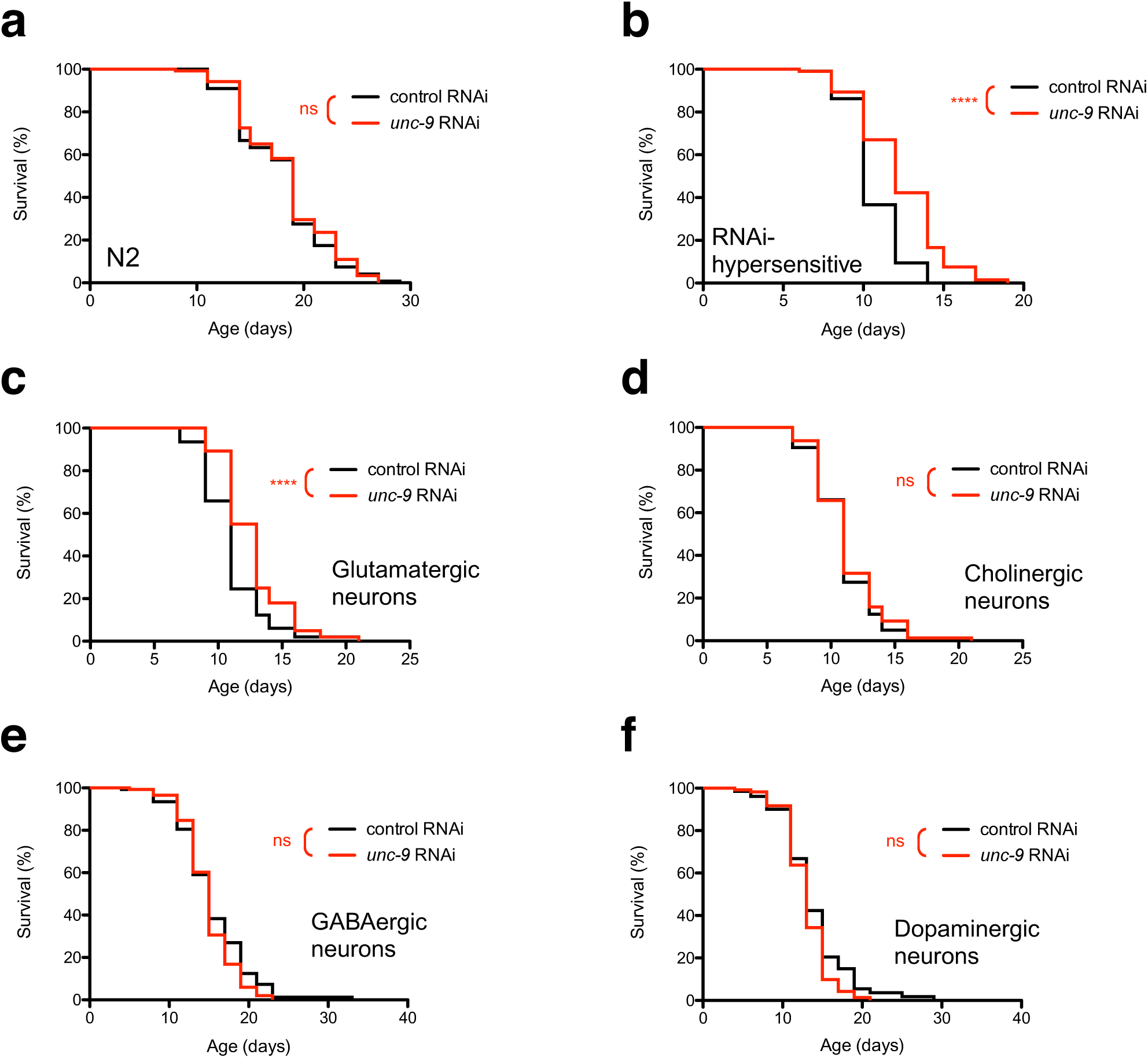
*unc-9* acts in the glutamatergic nervous system to regulate lifespan. **a** RNAi knockdown of *unc-9* does not affect lifespan in N2 animals (n=139 animals per group), but (**b**) increases it significantly in the RNAi hypersensitive KP3948 strain (N=130 animals per group). **c** Lifespan is significantly increased by RNAi knockdown of *unc-9* in glutamatergic neurons (N=130 animals control; 129 *unc-9* RNAi), but not in (**d**) cholinergic (N=130 animals per group), (**e**) GABAergic (N=136 animals control; 140 *unc-9* RNAi) or (**f**) dopaminergic neurons (N=134 animals control; 136 *unc-9* RNAi). ****P<0.0001; ns not significant, using Log-rank test.

*Unc-9* is expressed in 108 of 118 neuron classes^20^. We therefore sought to determine whether the role of *unc-9* in modulating ageing is widely distributed across the nervous system or can be narrowed down to a specific group of neurons. To test this, we specifically and selectively knocked down *unc-9* in four domains of the nervous system: the glutamatergic, GABAergic, dopaminergic and cholinergic neurons. To this end we used a tissue-specific RNAi approach, where only defined subsets of the nervous system are susceptible to RNAi by feeding^46^. We found that lifespan was significantly increased when *unc-9* was selectively knocked down in the glutamate-releasing neurons (Fig. 5c); no effect was seen from *unc-9* knockdown in either cholinergic, GABAergic or dopaminergic neurons (Fig. 5d,e,f). This finding tantalisingly suggests that the effect of UNC-9-containing gap junctions on lifespan may be caused by its coupling of specific neurons or neural circuits.

### The glutamatergic *unc-9*-expressing neurons are predominantly mechanosensory

To further narrow down the neurons where *unc-9* modulates ageing, we made use of the fact that the expression patterns of both *unc-9* and the vesicular glutamate transporter *eat-4* have been mapped to the level of individual neurons^20, 47, 48^. Their expression overlaps in 31 neuron classes of the somatic nervous system, namely the sensory neurons ADL, ALM, AQR, ASE, ASH, ASK, AVM, AWC, FLP, OLL, OLQ, PHA, PHB, PHC, PLM, PQR, PVD and URY; the interneurons ADA, AIB, AIM, AIZ, AUA, DVC, LUA, PVQ, PVR, RIA and RIG; the motoneuron RIM; and the polymodal IL1 sensory/inter/motoneurons (Fig. 6a; Supplementary Fig. 5). Additionally, they overlap in three pharyngeal neuron classes.

**Fig. 6.**
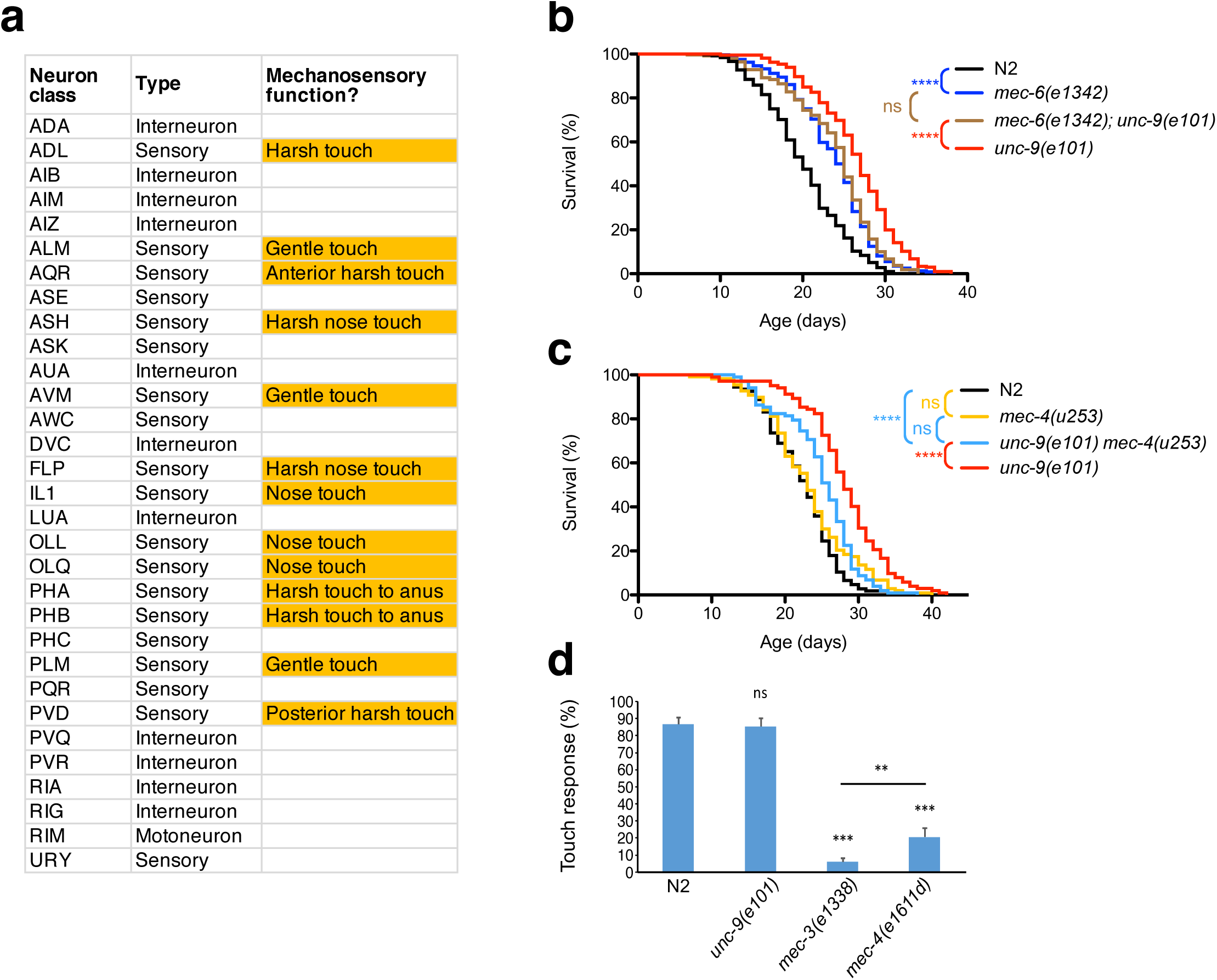
UNC-9 depends on mechanosensation to regulate lifespan. **a** Class, type and mechanosensory function of glutamatergic neurons expressing *unc-9*. See also Supplementary Fig. 5. **b** Lifespan of *mec-6(e1342); unc-9(e101)* mutants is identical to that of *mec-6(e1342)* mutants, significantly shorter than in *unc-9(e101)* mutants, but longer than in N2 controls (N=150 animals per group). **c** Lifespan of *unc-9(e101) mec-4(u253)* mutants is significantly shorter than in *unc-9(e101)* mutants, whereas *mec-4(u253)* mutants have similar lifespan to N2 controls (N=120 animals per group). **d** Touch responses of N2, *unc-9(e101)*, *mec-3(e1338)* and *mec-4(e1611d)* animals (N=15 animals per group). **P<0.01; ***p<0.001; ****p<0.0001; ns not significant; (**b**), (**c**) Log-rank test; (**d**) Student’s t-test.

These neuron classes show intriguing shared features: they are dominated by sensory neurons – 19 of the 31 classes are sensory neurons. Sensory perception plays a central role in regulating *C. elegans* lifespan^49, 50^, and chemosensory^51^, thermosensory^52, 53^ and oxygen-sensing neurons^54, 55^ have been shown to shorten or extend the animals’ lifespan in an environmental context-dependent manner. The oxygen-sensing neurons AQR and PQR signal tonically at 21% ambient oxygen but are largely inactive at reduced O_2_ levels (7%)^56^. If UNC-9 acted in those neurons to limit lifespan, we would therefore expect to see a difference in lifespan between animals grown at 21% or 7% O_2_. However, lifespan in both N2 and *unc-9* mutant animals did not significantly differ between animals grown at 21% and 7% oxygen (Supplementary Fig. 4). Of the above neurons which are chemosensory, ablating ASE, ASK and AWC individually does not affect lifespan^51^.

Strikingly, most of the glutamatergic *unc-9*-expressing sensory neurons (13 of 19) are implicated in mechanosensation, including both gentle and harsh touch to the head or body. They constitute a majority of the 20 known mechanosensory neuron classes in *C. elegans* (Fig. 5a; Supplementary Figure 5) ^57, 58^. Moreover, several of the glutamatergic interneurons that express *unc-9*, namely PVR, LUA, AIB and RIG, receive direct synaptic connections from the mechanosensors^59^. The mechanosensory and interneurons feature considerable electrical coupling both with each other and with the command interneuron circuit, particularly AVD and PVC, which control the direction of movement and also express *unc-9* (Supplementary Fig. 6).

### UNC-9 depends on mechanosensation to regulate lifespan

Our observations raise the possibility that the regulation of lifespan by UNC-9 gap junctions depends on a neural circuit involving mechanosensory neurons. We hypothesised that in that case the increased longevity conferred by the lack of *unc-9* would be affected if worms are insensitive to touch and consequently the mechanosensory circuit less active. Mechanosensation has not hitherto been known to affect *C. elegans* lifespan

To test this hypothesis, we investigated if there is a genetic interaction between the *unc-9* loss-of-function mutation and mutants of genes required for touch sensation, namely of the degenerin mechanosensory channel subunit MEC-4 and of the paraoxonase-like protein MEC-6, which are expressed in the touch receptor neurons ALM, AVM, PLM and PVM. Null mutants of these genes are defective in responding to gentle touch and eliminate both mechanoreceptor currents and touch-evoked Ca^2+^ transients^60, 61^. We performed lifespan assays with double mutants of *mec-6; unc-9* and *unc-9 mec-4* and compared them with single mutants of either gene as well as with N2 wild-type. Both double mutants were defective in touch responses in the same way as *mec-6* and *mec-4* single mutants are (Supplementary Fig. 7). We found that *mec-6* null mutants live longer than N2 but expand median lifespan less than *unc-9* null mutants (5 vs. 7 days) (Fig. 6b; Supplementary Table 2). Lifespan of the *mec-6; unc-9* double mutants was identical to that of *mec-6* single mutants but longer than in N2 and shorter than in *unc-9* animals. The longevity conferred by the absence of *mec-6* therefore is epistatic to *unc-9* and does not depend on this innexin; conversely, loss of *unc-9* depends on a functional *mec-6* gene to prolong life.

*Unc*-9 also depends on *mec-4* to expand lifespan. Longevity of the *unc-9 mec-4* double mutant was significantly shorter than in the *unc-9* mutant alone, while lifespan of *mec-4* mutants was not significantly different from that of N2 animals (Fig. 6b,c). We conclude that the increase of lifespan in *unc-9* mutants depends on touch sensation.

We then sought to test if *unc-9* mutants themselves are defective in gentle touch sensation and found that these mutants had wild-type responses to touch, indicating that the increase in lifespan of *unc-9* mutants is not caused by decreased mechanosensation (Fig. 6d). To further determine which neurons in the mechanosensory circuit play a role in the control of ageing by UNC-9, we knocked down *unc-9* in the touch receptor neurons by RNAi.

Separately, we ablated the PVR interneuron, which is heavily synaptically connected with mechanosensory neurons, using a laser microbeam. However, these treatments did not affect lifespan (Supplementary Fig. 8), suggesting that in addition to the touch-receptor neurons, other glutamatergic neurons such as harsh touch sensors may be involved in the regulation of longevity by UNC-9.

### UNC-9 alters age-dependent morphological changes of the touch receptor neurons

In both humans and *C. elegans*, healthy ageing of the nervous system is marked primarily by a decline in the structural integrity of neurons with age rather than outright neuron loss^62^. These age-associated morphological changes are particularly prevalent in the touch-sensing neurons of *C. elegans* and correlate with decreased response to light touch as well as decreased mobility^63^. Because *unc-9* mutants depend on a functional mechanosensory machinery for their positive effect on lifespan, we speculated that the absence of this innexin may alter the progressive decline of touch receptor neuron integrity with age. To test this, we imaged and assessed the morphology of the gentle touch-sensing ALM, AVM and PLM neurons in ageing worms. The cell bodies of ALM and AVM are positioned anteriorly, while those of PLM are located posteriorly (Fig. 7a). In young adults, the cell bodies are typically round or oval but develop progressively more irregular shapes with age^64^. All three extend a major neurite anteriorly which make synaptic contacts with other neurons. In addition, ALM and PLM form short posterior processes^65^. In young animals, those processes appear straight and uniform but develop various abnormalities in old adults, namely a wavy appearance and protrusions of different shapes (Fig. 7b)^66, 67^. We found that *unc-9(101)* mutants altered the age-dependent decline in morphological integrity, and had a markedly different effect in different cell types. In the ALM neurite, the progressive increase in abnormalities was significantly accelerated in *unc-9* mutants, while degeneration in the PLM dendrite was slower than in N2 (Fig. 7c,d). In AVM, cell bodies showed a significantly faster deterioration in the *unc-9* mutants (Fig. 7e). Integrity of the ALM or PLM cell bodies was not significantly affected by genotype (Supplementary Fig. 9a,b).

**Fig. 7.**
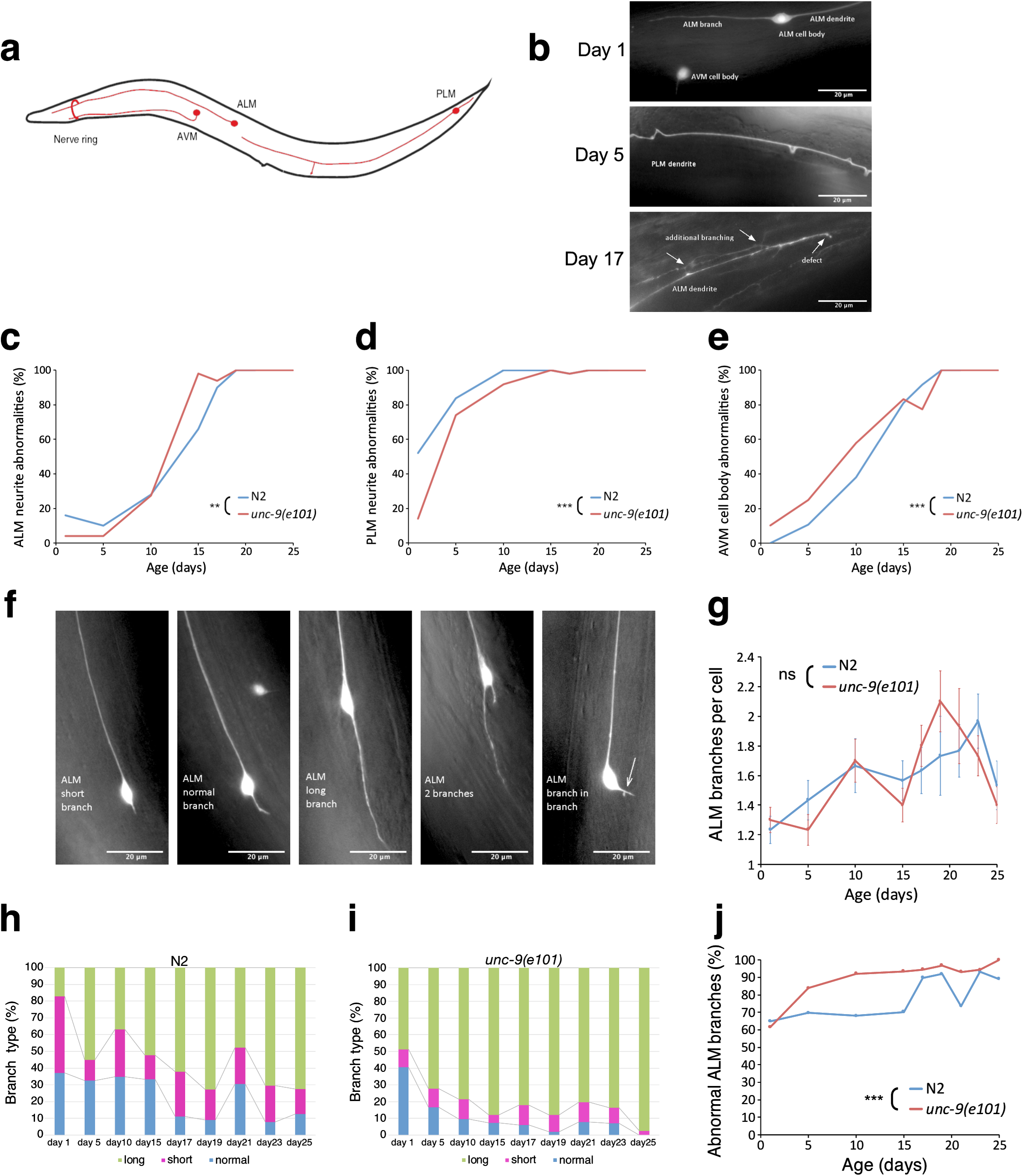
UNC-9 alters age-dependent morphological changes of the touch receptor neurons. **a** Schematic representation of ALM, AVM and PLM morphology. **b** Representative GFP images of mechanosensory neurons in day 1, day 5 and day 17 *unc-9(e101)* adults; arrows showcase ageing-related defects; scale bar: 20 µm. **c** Age-related degeneration of ALM and (**d**) PLM neurites, and of (**e**) AVM cell body in *unc-9(e101)* mutants and N2 controls. **f** Example GFP images of normal branching and age-related morphological defects arising in ALM neurites; scale bar: 20 µm. **g** The overall number of ALM branches increases similarly with age in *unc-9(e101)* mutants and N2 controls. **h** The increase in proportion of long branches with age, relative to short and normal branches combined, is significantly lower in N2 (**i**) than in *unc-9(e101)* animals (p<0.001, ***). **j** Abnormal branching rises earlier in *unc-9(e101)* mutants than N2 controls. **P<0.01; ***P<0.001; ns not significant; (**c-e**), (**j**) binomial regression model; (**g**) Poisson generalised linear model; (**h,i**) multinomial regression model. N=50 animals per group for (**c-e**); N=30 animals per group for (**g-j**).

**Fig. 8.**
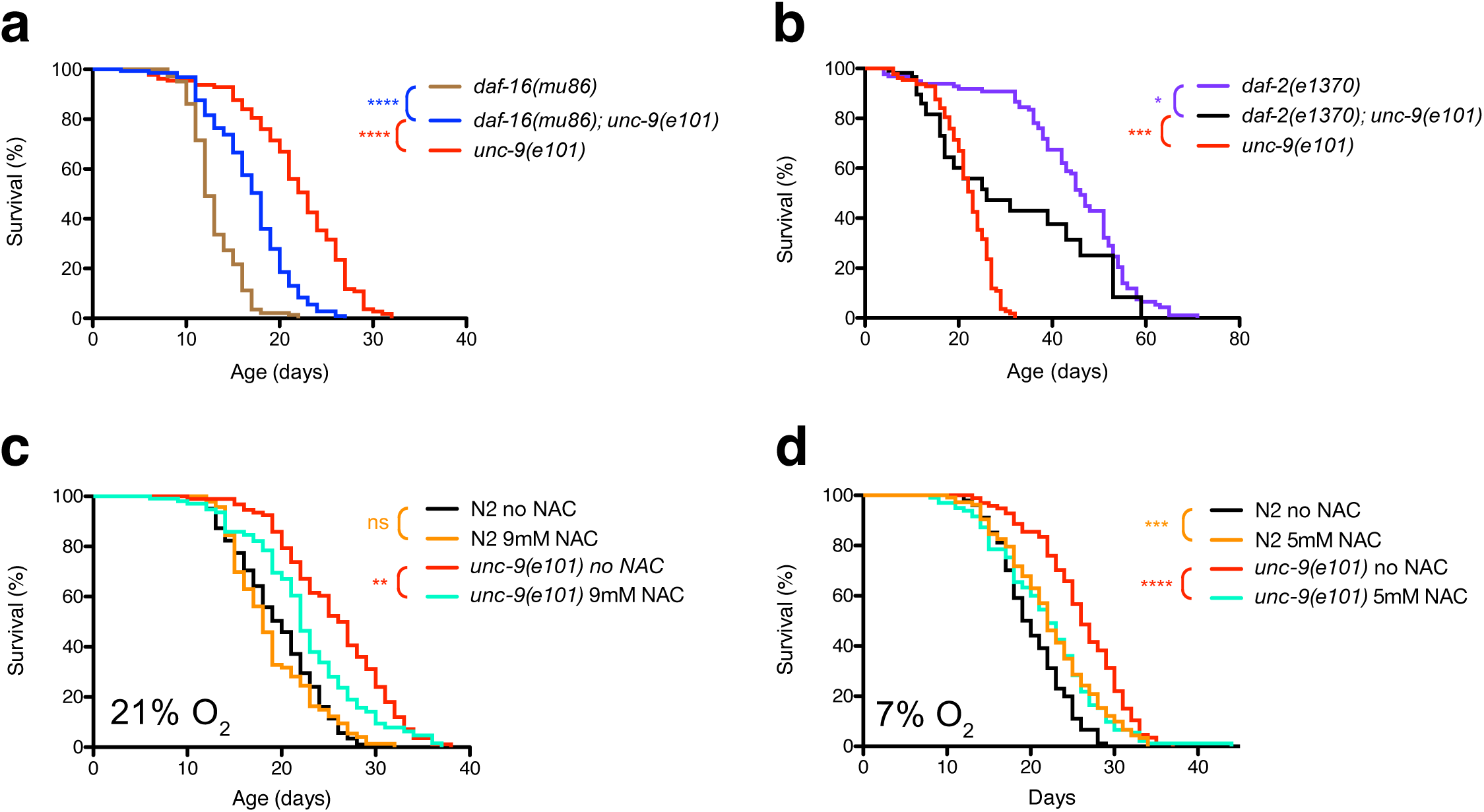
UNC-9 lifespan regulation is independent of FOXO signalling but linked to reactive oxygen species. **a** Lifespan of *daf-16(mu86)*, *daf-16(mu86); unc-9(e101)* and *unc-9(e101)* mutants. **b** Lifespan of *daf-2(e1370)*, *daf-2(e1370); unc-9(e101)* and *unc-9(e101)* mutants. **c** Lifespan of *unc-9(e101)* mutants and N2 animals with or without treatment with 9mM N-Acetyl-Cysteine (NAC) maintained at normoxia (21% O_2_). **d** Lifespan of *unc-9(e101)* mutants and N2 animals maintained at hypoxia (7% O_2_) with or without treatment with 5mM NAC. *P<0.05; **p<0.01; ***p<0.001; ****p<0.0001; ns not significant, using Log-rank test. N=150 animals per group (**a**),(**b**); N=120 animals per group (**c**),(**d**).

We conclude that *unc-9* affects the progression of neuronal ageing in a markedly different way between touch neuron subtypes, where specific morphological structures age faster, are unaffected or deteriorate slower in the mutant. This finding parallels previous reports showing that age-dependent morphological degradation in the touch receptor neurons differs markedly between neuronal subtypes^66, 67^.

Another hallmark of touch receptor neuron degeneration is the appearance of additional branches emanating from either the somata or processes in ageing worms^63, 67^. In the above experiment, we observed that the touch receptor neurons in older *unc-9* mutants appeared to have longer and more abnormal branches than N2 animals (Fig. 7f). We therefore decided to quantify the morphology of neuronal branches of ageing worms, focusing on the ALM neurons where it could be observed most clearly. We found that the overall number of branches in ALM progressively increases with age, but is not affected by the *unc-9* genotype (Fig. 7g). On the other hand, the fraction of long branches (≥2x soma diameter) was significantly increased in *unc-9* (Fig. 7h,i). This was already apparent in young adults but increased further with age. The proportion of all types of abnormal branches combined - comprising short or long branches, loops and branching out of existing branches - increased with age in both genotypes but rose much earlier in unc-9 mutants (Fig. 7j).

Taken together, these results show that loss of *unc-9* alters the morphological ageing of the touch receptor neurons. Notably, *unc-9* mutants have an accelerated appearance of ageing hallmarks in the ALM and AVM neurons.

### UNC-9 lifespan regulation is independent of FOXO signalling but linked to ROS

Our results point to a novel role of gap junction coupling in mechanosensory circuits in regulating *C. elegans* ageing. This raises the question which ageing pathways are implicated in the effect of *unc-9* on lifespan. Insulin/insulin-like growth factor-1 factor signalling (IIS) plays a pivotal role in the regulation of longevity^68^. In *C. elegans*, activation of the insulin receptor DAF-2 stops DAF-16, the *C. elegans* orthologue of the FOXO transcription factor, from entering the nucleus. When DAF-2 is inhibited, DAF-16 translocates to the nucleus and activates genes involved in stress resistance and longevity. Consequently, *daf-2(e1370)* mutants with reduced IIS signalling live twice as long as N2 wild type, while *daf-16(mu86)* null mutants have shortened lifespan. To test if *unc-9* acts through IIS signalling to regulate lifespan, we created double mutants of *unc-9(e101)* with both mutants. We found that *daf-16; unc-9* animals lived significantly longer than *daf-16* single mutants (Fig. 7a). Median lifespan of the double mutants is 50% longer than in *daf-16* alone (18 vs. 12 days). This indicates that *unc-9* null mutants increase longevity independently of *daf-16/*FOXO signalling. The lifespan of the *daf-2; unc-9* double mutant population appeared to fall in two categories: for those dying before they reach median lifespan of the population, it was the same as in *unc-9* single mutants; in contrast, those who survived beyond this age lived considerably longer than *unc-9* mutants. This suggests that *unc-9* is partially required for the lifespan increase conferred by reduced IIS signalling. However, the analysis of the *daf-2; unc-9* double mutants was impeded by their strongly increased propensity for internal egg hatching, which eventually kills the parent, with one event observed as late as day 51 (Supplementary Fig. 10); we presume that this is caused by a combination of an increased rate of internal hatching caused by the *unc-9* mutation and the ability of *daf-2* mutants to produce progeny late in life^69^.

Together, we conclude that *unc-9* acts independently of *daf-16*/FOXO to modulate ageing, while our data do not rule out an interaction with *daf-2* insulin receptor signalling.

Intercellular gap junction channels enable the diffusion of small molecules from cell to cell. It is well established that this can cause a “bystander” effect where cells experiencing oxidative insults or apoptotic cell death increase the damage or death of gap junction-coupled cells through the diffusion of as yet unknown signals^12, 70^. We therefore aimed to explore if oxidative stress may be implicated in the life extension by *unc-9* mutants. We thus conducted a lifespan assay where animals were grown in the presence of N-Acetyl-Cysteine (NAC), an antioxidant and scavenger of all types of reactive oxygen species (ROS)^71^.

N2 worms maintained at normoxia (21% O_2_) did not respond to NAC treatment, consistent with previous reports^72, 73^. In contrast, 9mM NAC eliminated a substantial part of the life extension conferred by the loss of unc-9, with median lifespan increasing by only 2 days instead of 6 in untreated *unc-9* mutants, relative to wild-type (Fig. 7c).

An important consideration in how ROS levels govern longevity is that the natural habitat of *C. elegans* contains much lower levels of oxygen than the laboratory conditions under which most experiments are conducted^56, 74^. We therefore also maintained strains at their preferred and more physiological oxygen tension of 7% O_2_^56^. At this lower oxygen tension, N2 wild-type animals survive significantly longer when grown with NAC (Fig. 7d). The effect of NAC on lifespan appears to be mostly affecting the survival of older animals. In contrast, NAC treatment again strongly reduced the increased lifespan of *unc-9* null mutants (Fig. 7d). A higher dose of 9mM NAC had similar but weaker effects on worms maintained at 7% O_2_ (Supplementary Fig. 11a,b).

Together, these results indicate that the increased longevity conferred by loss of the *unc-9* innexin substantially requires ROS and that antioxidant activity can increase longevity in wild-type but not *unc-9* mutant animals.

## Discussion

Gap junctions are unique and ubiquitous intercellular conduits that provide electrical, metabolic and biochemical coupling between cells^5, 6^. They also play key pathological roles such as the propagation of cellular injury or death signals to “bystander” cells^10–12^. Despite their fundamental importance in intercellular communication, it has not been known whether gap junctions and the genes forming them contribute to the regulation of ageing and longevity.

We now show that the genes encoding gap junction subunits have diverse and widespread roles in regulating *C. elegans* ageing. A recurrent difficulty in studying gap junction genes arises from their partial redundancy, where members of this gene family have overlapping roles and thus when one gene is missing, others can compensate for its absence^75^. We found that despite their frequent overlap in expression, the majority of individual innexin mutants modulate lifespan, and thus have non-redundant effects. This suggest that the modulation of lifespan is specific to individual innexins acting in defined sets of cells, rather than a universal feature common to all of them. It does not rule out that innexins have overlapping roles and thus when one innexin gene is missing, others can compensate for its absence. Future studies may examine if double mutants of innexins with similar expression patterns, particularly in the nervous system, show a further increase in lifespan.

A major finding of this study is that loss of the *unc-9* innexin increases lifespan. Our results suggest that the presence of UNC-9 in the glutamatergic nervous system limits longevity. It acts in the same pathway as UNC-7 and requires regulation by the stomatin UNC-1 to control lifespan.

The effect of UNC-9 on lifespan could potentially be caused by a nonjunctional function of the protein. However, we found *unc-1* to regulate lifespan by acting on *unc-9*. *Unc-1* encodes a stomatin-like integral membrane protein that colocalises with UNC-9 and specifically modulates the gating of UNC-9 gap junction channels in muscle, but is not required for the expression or subcellular localisation of UNC-9^76^. In the absence of UNC-1, UNC-9-containing gap junctions are thought to be mainly in the closed state^76, 77^. Our results support the hypothesis that the role of UNC-9 in controlling longevity rests on its function as an intercellular channel and that its open state causes a shortening of *C. elegans* lifespan.

Some studies reported that lifespan-extending mutations predominantly increase the frail period, while other reports concluded that mutants such as *daf-2* do increase healthy ageing^39, 40, 43, 78–80^. We show here that *unc-9* mutants enjoy not only an increase in lifespan but also extend the period of healthy ageing, with locomotory activity, responsiveness, defecation and pharyngeal pumping all persisting longer with age than in wild type. However, *unc-9* mutants in general bear multiple phenotypic defects, especially uncoordinated behaviour. Our results suggest that by selectively removing *unc-9* from glutamatergic neurons, it would be possible for worms to enjoy the benefit of increased longevity without having other behavioural defects.

*unc-9* is expressed in nearly all neurons, potentially creating an electrically coupled network encompassing virtually the entire nervous system, and is also expressed in most muscles. Yet we found that it controls lifespan specifically in a subset of its expression domain, namely the glutamatergic neurons. This raises the possibility that a functional circuit consisting of these neurons regulates lifespan in an UNC-9 gap junction-dependent manner. While we cannot rule out that other *unc-9*-expressing cells also have a role in lifespan regulation by this innexin, our evidence argues against it. Although expressed in the pharyngeal nervous system, *unc-9* were not feeding deficient and had only a slight reduction in brood size. It is therefore unlikely that the increase of *unc-9* mutant lifespan is caused by dietary restriction or reduced reproduction. It is also unlikely that the uncoordinated phenotype and reduced locomotory activity of *unc-9* mutants causes their lifespan extension. The *unc* phenotype of *unc-9* is thought to be caused by dysfunction of the locomotory nervous system, not of the largely sensory glutamatergic neurons. Moreover, previous reports^81, 82^ have found that most *unc* mutants tested do not increase lifespan.

*Unc-9* is expressed in glutamatergic sensory neurons that respond to chemical, temperature and oxygen cues which are known to influence lifespan. However, these neurons do so by regulating *daf-16* FOXO signalling^51, 54, 83^. Our data show that *unc-9* acts independently of *daf-16* to regulate longevity, making it unlikely that *unc-9* regulates lifespan in these neurons. Instead, our findings point towards a very unexpected set of neurons involved, namely the mechanosensory neurons. These had not previously been implicated in lifespan regulation. However, studies have shown that mechanosensory neurons are particularly prone to decline of morphological integrity with age^63, 66^. Changes in the mechanical properties of cells are hallmarks of the ageing process, which correlates with progressive decline in the structural integrity of cells and their diminished response to mechanical forces^84^. It is conceivable that mechanosensory cells represent a link between mechanical stress, the age-dependent decline of mechanical properties, and the mechanisms that regulate organismal ageing. In that respect, it is interesting that several mammalian connexins have been shown to be mechanosensitive^85^, and investigating how their properties change with age could be the focus of future studies.

We observed that loss of *unc-9* alters the structural integrity of the touch receptor neurons, and according to several measured parameters can accelerate the deterioration of their morphology. This suggests that isolation of these neurons from electrical neuronal networks worsens their state, but paradoxically improves lifespan and healthspan of the whole organism.

In that regard, it is interesting that ROS are partially required for the long life of *unc-9* null mutants. This suggests that the mechanism of how gap junctions impinge on ageing is linked to either redox regulation or oxidative stress, key factors in both cellular ageing and longevity^86^. The nature of this mechanism remains to be uncovered by future studies. In any electrically coupled neuronal circuit, cells will be in different activity states, resulting in a current flow from more active to less active neurons. It has been shown that such shunting can regulate the function of a mechanosensory circuit in *C. elegans*, where chronically inactive electrically coupled neurons inhibited sensory responses in the whole circuit^87^. Our observations could potentially be explained by a model where, similarly to neuronal injury models, noxious or ageing-promoting signals might be spreading across *unc-9*-containing gap junction channels to connected neurons, for example reactive byproducts originating from neurons or neural circuits with strong, persistent neural activity to “bystander” neurons with lower activity, who thereby are damaged or functionally impaired after long-term exposure. Alternatively, the absence of *unc-9* gap junctions may alter cellular metabolism and thus their redox state as a result of glutamatergic cells being isolated from other neurons.

Of the other innexin mutants we found to increase lifespan, *inx-15* is particularly interesting as it had the second strongest effect. It is only expressed in the intestinal epithelial cells and has not been characterised previously. The intestine is a key player in the control of lifespan and develops multiple pathologies with age^88^. Old *C*. *elegans* have reduced immunity and diminished capacity to control their intestinal bacterial accumulation^89^. Bacteria, the worms’ exclusive food, can colonise their guts and invade their tissues. In mammals, connexins in the gut can play a disease-promoting role and spread bacterial infections and inflammation across cellular networks. It will be interesting to see if *inx-15* plays a role in the interaction between intestinal cells and gut bacteria.

The innexins expressed in the germline had opposing effects – *inx-14* mutants extended lifespan, while *inx-21* and *inx-22* reduced it. *Inx-14* is required for both germ cell proliferation and negatively regulates oocyte meiotic maturation, while inx-21 contributes only to germ cell proliferation and inx-22 only to inhibiting meiotic maturation^90^. It may be that removal of both functions resembles removal of the germline, which prolongs lifespan, but that inhibiting each process individually reduces longevity.

While cellular mechanisms of ageing have been the focus of intense investigation, it remains much less well understood how non-cell-autonomous mechanisms provide a systemic control of ageing and longevity. The nervous system acts via diverse pathways to modulate organismal ageing and lifespan and itself undergoes age-dependent deterioration of its function and structure^91, 92^. A full understanding of the causative relationships between age-dependent decline of cells in the brain, organismal aging and longevity remains elusive. Our study offers the tantalising prospect that gap junction intercellular communication underpins a previously unknown non-cell autonomous mechanism that modulates ageing.

## Methods

### Nematode strains

*C. elegans* strains used are listed in Supplementary Table 3. All strains were maintained under standard conditions^93^ in 5 cm Petri dishes on NGM (nematode growth medium) agar seeded with *E. coli* OP50. Except for *unc-1(e1598)*, which is a gain-of-function mutation, all other mutants used are assumed to be loss-of-function mutants. The precise molecular lesion in the *inx-5(ok1053)* and the *inx-18(ok2454)* mutants had not been previously determined; both were sequenced and found to bear a 1221 bp deletion in *inx-5* (nt 5058-6278 in R09F10) and a 1704 bp deletion in *inx-18* (nt 30705-32408 in C18H7), that in both cases remove more than one exon and are therefore putative null mutants (Supplementary Table 1). The *inx-6(rr5)* loss of function allele is temperature-sensitive and is fully restrictive at 25°C^1^; it was tested at 20°C to allow cross-comparison with other innexin mutant strains in the assay.

*Unc-9* rescue lines were generated by microinjection^94^ of a DNA Mix containing 1 ng/µl of *unc-9* encoding fosmid WRM0611aH10 and 1.5 ng/µl of pCFJ90 (pmyo-2::mCherry) into *unc-9(e101)*. After 2 days F1 animals were singled by co-injection marker selection. Stable arrays were identified in the transgenic F2 generation.

### Lifespan assays

For lifespan assays, established protocols were followed, with some adaptations^49, 95, 96^. Strains were first cultured under standard growth conditions for two generations to minimise non-genetic effects on lifespan. Then, 30 third-generation 2-day old adult hermaphrodites were allowed 4-8 hours for timed egg-laying and progeny laid during this time was grown for approximately two days at 20 °C to late L4 larval stage, before being transferred to fresh plates at 12-15 animals per plate. For each strain, animals were split across ten plates. Animals were passaged to fresh plates every second day during the reproductively active stage (10 days) and then passaged every three days until the end of the experiment, or when a contamination occurred. Each plate was scored every day for live animals, dead animals and censored animals. Lifespan was measured from day 1 of adulthood and all lifespan assays were conducted at 20°C. To establish if an older worm in stationary state was dead or alive, a worm pick or a human eyelash mounted on a pick was used to gently brush the animal on different parts of the body, namely, the tail, the ventral area and the nose. If the animal did not react after several attempts, then it was considered dead and was removed. Worms were censored when they escaped from the plate, if they became contaminated by bacteria or fungi, displayed internal egg hatching, vulva rupture or vulva protrusion, or were mishandled^95, 96^.

### Lifespan analysis

Analyses of lifespan data were conducted using a Kaplan-Meier survival analysis on GraphPad Prism versions 6 and 7. For statistical comparisons two-tailed Log-rank tests were performed. Median Survival represents the time at which half the animals in a given experiment have died. Along with median survival, which represents the central tendency of survival scores, 25th and 75th percentiles and the inter-quartile range (IQR) were reported as a measure of dispersion. Most survival score distributions observed were multimodal, which means that the arithmetic mean is not informative. Also, in contrast to the 25th and 75th percentiles, minimum and maximum lifespan are very vulnerable to outliers and therefore not reported. Median survival and IQR are also represented by a white dot and a black box, respectively, in the Violin plots (Supplementary Fig. 1) along with the probability density of the data at different values, smoothed by a kernel density estimator (KDE). Violin plots are a combination of box and density plots and allow a better understanding of how the scores are distributed, which is especially useful for asymmetric distributions. All Violin Plots were generated with Seaborn’s violin plot function in Python 3.7.

### RNAi by feeding

RNAi by feeding was performed as described previously^46, 97^ with a few adaptations for the lifespan assays. Standard NGM plates were supplemented with 25 μg/ml carbenicillin and 1 mM isopropyl b-D-1-thiogalactopyranoside (IPTG). The plates were allowed to dry in a dark environment at room temperature for five days before use. The RNAi clones containing the *oac-47* pseudogene (T22H2.2) and *unc-9* (R12H7.7) are from the Ahringer RNAi feeding library and were initially streaked on LB plates containing 25 μg/ml carbenicillin and 10 μg/ml tetracycline and grown overnight at 37°C. The next day, cultures were inoculated in 2x YT medium containing 25 μg/ml carbenicillin overnight at 37°C in a shaking incubator. Two drops from the inoculated culture were seeded onto each plate. Animals were transferred to the RNAi bacteria in L4 stage. *oac-47*, a pseudogene, served as negative control. Identity of the clones was confirmed by sequencing prior to use. Lifespan assays were conducted as described above except that animals were scored every other day.

### Locomotion assay

Measurement of locomotory speed decline was performed as described in ^98^ with some adaptations. 45 synchronised hermaphrodites from each strain (N2, CB101) where recorded for 5 min from day 1 of adulthood and every other day until death. Worms were transferred to fresh plates, seeded the day before, 15 min before the recording and the lid was removed for the last 2 minutes before recording, allowing adaptation to the air flow. The function Video Capture of the Wormlab setup (MBF Bioscience) and a Basler acA2440 camera were used. The selected frame rate was 14 fps, the video mode was 1600*1200 and the rest of the settings were used as pre-set from manufacturer. Analysis was performed using the tracking function of the software and moving average speed (µm/s), averaged across 7 frames, was extracted for each video. The values provided were further analysed in Excel. The absolute values of the results provided by the software were used for averaging one single value per video. Due to the lower initial locomotory speed of *unc-9* mutants, speed was normalised relative to the maximum locomotory speed observed in day 3 adults.

### Responsiveness assay

To assess responsiveness to light touch with a worm pick, we performed a lifespan assay as described above and evaluated touch responses every other day. To score response to touch we used an index where a score of 4 indicated that upon plate tap, animals initiated an escape response, typically a reversal, changed their direction of movement, or moved their nose; 3 if they responded in the same way to light touch to their tail; 2 if they responded to light touch along the body; and 1 if they responded to light touch to the nose. Animals that did not respond at all were considered dead and therefore not scored. Each worm was given a score which were then averaged across 12 animals to give a plate score for each of the 10 plates per strain.

### Defecation assay

The protocol to measure defecation cycle length was adapted from^44, 99^. Hermaphrodites were synchronised by timed egg laying and maintained at 20°C, on standard NGM plates, in the same as for lifespan assays. The defecation cycle length was measured on day 1 of adulthood and every third day onwards until expulsions were no longer visible. Animals were transferred to fresh plates and were allowed to adapt for at least 15 min before observing. Measurements were obtained from feeding, non-roaming animals and the duration of the cycle was defined as the time from one expulsion to the next. Three defecation cycles were measured for each animal.

### Pharyngeal pumping assay

Worms were examined *in situ* on NGM agar plates using a Leica S6D Greenough stereomicroscope with a Leica KL300 LED light source and a Point Grey Grasshopper3 USB3 camera, attached via a Leica 0.63x relay lens. Films were captured for 60s at 25fps and 2448×2048 resolution using FlyCapture 2.13.3.31 (Point Grey) at maximum zoom and encoded using M-JPEG compression with a quality ratio of 100 (maximum). When necessary, plates were moved gently to keep the head of the animal in focus and within the field of view. The number of pharyngeal pumps was scored manually by two independent observers for each worm by playing back the videos with VLC Media Player at 0.12x (day 2) to 1.00x (day 18+) speed and using a clicker counter.

### Brood size analysis

Brood size was measured by transferring single L4 worms to an NGM plate and kept at 20°C. Each adult worm was transferred to a fresh plate every day and the previous plate was kept at 20 °C for another 2 days, when the number of alive progeny was scored. This procedure was repeated until no alive progeny were found anymore.

### Gentle touch sensation assay

Experiments were performed on day 1 adults. Animals were placed on fresh 5 cm NGM plates seeded two days before with a layer of OP50 that did not touch the edges of the plate. After 10 min of acclimatisation, touch sensation was tested by streaking animals gently at the side with an eyelash pick^100^. In particular, a total of 10 stimuli per animal were given by alternating anterior (2^nd^ sixth of body) and posterior (5^th^ sixth of body) strokes every 3-4 sec. Positive reactions were defined as movement in the opposite direction of stimulus or stop of present movement^101, 102^. Average gentle touch responses of strains were compared. For statistical analysis unpaired Student’s t-test was used.

### Scoring of mechanosensory neuron morphology

To record neurodegeneration from the mechanosensory neurons, we followed published protocols^66, 67^. Briefly, for synchronisation, 600 L4 larvae per strain were picked. After reaching day 2 adulthood they were used for egg-laying. 30 animals per 50 mm NGM plate, seeded with OP50 bacteria, were allowed to lay eggs for 4-6 hours. Afterwards adults were removed and plates were kept at 20°C until the progeny reached L4 larval state. 60 L4s per strain were transferred to 25 NGM plates, seeded with OP50 bacteria, to reach a total of 1500 animals per strain. Animals were transferred to fresh plates every second day to remove progeny.

Scoring was performed on days 1, 5, 10, 15, 17, 19, 21 and 25 on 50 animals per strain and day. For scoring, 10 animals were placed on a pad with 2% agarose in M9, immobilised with 50 mM levamisole, covered with a coverslip and mounted on a Zeiss Axio Imager.M2 upright fluorescence microscope. Neurons were scored by eye at 40x magnification. In each animal, one ALM neuron (cell body and dendrite), AVM (cell body), and PLM (one dendrite, left and right cell body), were scored and classified as intact, defective or degenerated. Afterwards example images were taken with ZEN pro imaging software. For analysis, the average of classifications per strain, neuron, neuron-part and day was compared.

The scoring of ALM branching was performed in the same way by collecting data and branch categories were defined as follows: 1-2x diameter soma = normal branch, less than 1x diameter soma = short branch, 2x or more diameter soma = long branch, secondary branch emanating from an existing branch out of the soma = branch in branch, branch grows in a loop = loop. From this information the number of branches out of cell body, number of branches in total, normal vs. abnormal branching and the respective branch types per strain and day were compared.

### Statistical analysis

Data were analysed using SPSS 24 Statistics software (IBM). Normality was assessed using the Shapiro-Wilk test. Unbalanced data with normal distributions were analysed using linear mixed models with type III test of fixed effects. Non-normal data showing heteroscedasticity under Levene’s test were analysed using a generalised estimating equations generalisation with Wald’s test for linear models. Binomially distributed data were analysed using a binomial logistic regression model. Multinomially distributed data were analysed with a multinomial logistic regression model. Data where the response variable was a count were analysed using Poisson regression.

## Acknowledgements

We thank the Caenorhabditis Genetics Center and the National Bioresource Project for the nematode for strains and Maria Doitsidou and Sebastian Greiss for helpful suggestions. We also thank Simon Warburton-Pitt for help with genotyping, Kanyarat Benjasupawan for contributing to healthspan measurements, Vishaka Kulkarni for help with lifespan assays, and Eugenia Goya for advice on scoring neurodegeneration. We gratefully acknowledge financial support by the Wellcome Trust (109614/Z/15/Z), the Medical Research Council (MR/N004574/1), the Muir Maxwell Epilepsy Centre and the University of Edinburgh (PhD studentship to NAV).

## Author contributions

K.E.B., N.A.V. and K.E.F. conceived and planned the study. N.A.V., M.N. and K.E.B designed the lifespan experiments which were performed by all authors and analysed by N.A.V., K.E.B. and K.E.F. Healthspan assays were planned, performed and analysed by N.A.V., E.D., D.C.M. and K.E.B. with help from A.A. Touch receptor experiments were designed, performed and analysed by K.E.F., D.C.M. and E.D. Crosses, genotyping and creating transgenic strains were done by K.E.F. with help from P.G. All authors discussed the data; and K.E.B. wrote the manuscript with input from N.A.V., K.E.F., E.D., D.C.M. and Q.L.

## Supplementary Methods

### Ablation

Laser ablation of the PVR interneuron to conduct a lifespan assay was performed essentially as described^16^. L1 larvae were immobilized in a 2 µl drop of 80 mM NaN_3_ on a 2% agarose pad. PVR was ablated in flp-20::GFP worms. PVR was identified based on its position near the PLM neurons and the shape of its processes. Worms were mounted on a Zeiss Axio Imager.M2 using a 63x objective, and cells were ablated with the Andor Micropoint system using a laser microbeam in asynchronous FRAP mode with 60 pulses at 60% of maximal power. The next day L4 stage worms were checked for proper ablation and selected for the lifespan assay. Mock ablated animals were treated in the same way except that the laser firing was omitted.

### Tap assay

Conditions and software were used as outlined for the locomotion assay, with the addition of mechanical stimuli. The tapper component of WormLab (MBF Biosciences) induces a single mechanical stimulus, and consists of a custom built linear solenoid actuator which drives a small plunger into a contact point on the plate holder assembly. The linear solenoid has a 5.5mm throw and is operated at 24V DC. Maximum stimulus intensity was used every 60 seconds during the recording. Analysis was performed as described for the locomotion assays. In addition, moving average speed was calculated before and after two taps. The subtracted value between the two time periods, indicating the change in speed, was averaged between the two taps and plotted as relative increase in speed^17^.

### Swimming assays

To verify the rescue effect of *unc-9* expression from a fosmid, the swimming behaviour was compared between N2 Bristol, *unc-9(e101)* and *unc-9(e101); [punc-9::unc-9; pmyo-2::mCherry]* worms. One day prior to the assay L4 larvae were selected. Ten animals per strain were transferred with an eyelash pick to one well of a 96 well plate, filled with NGM and topped up with M9 Buffer. After ten minutes of acclimatisation videos of 1 min length at 30 fps were recorded with a Point Grey Grasshopper3 camera mounted on a Leica stereo microscope. Thrashes, defined as the synchronous left and right movement of head and tail back to start position, were counted for each animal. Average thrash rates of strains were compared. For statistical analysis Student’s t-test was used.

### Crosses

#### unc-9(e101)/daf-2(e1370)

Because of temperature sensitivity of daf-2(e1370), crosses were performed at 15°C. The unc-9 mutation was verified by locomotion defective phenotype as well as PCR and restriction digest with BslI; the daf-2 mutation confirmed by PCR with primer pair KEB836/KEB837, restriction digest with BslI and sequencing.

#### unc-9(e101)/daf-16(mu86)

The unc-9 mutation was verified by locomotion defective phenotype as well as PCR and restriction digest with BslI; the daf-16(mu86) deletion mutation was confirmed by PCR with primer pairs KEB839/KEB840 and KEB838/KEB840.

#### unc-9(e101)/mec-6(e1342)

The unc-9 mutation was verified by locomotion defective phenotype as well as PCR and restriction digest with BslI; the mec-6(e1342) substitution mutation preselected by touch response and verified by sequencing of a PCR product generated with primer pair KEB914/KEB915.

#### unc-9(e101)/mec-4(u235)

The unc-9 mutation was verified by locomotion defective phenotype as well as PCR and restriction digest with BslI; the mec-4(u235) deletion mutation preselected by touch response and verified by PCR with primer pairs KEB911/KEB913 and KEB912/KEB913.

#### unc-9(e101)/pmec-3::GFP

The unc-9 mutation was verified by locomotion defective phenotype as well as PCR and restriction digest with BslI; presence of pmec-3::GFP was confirmed by GFP fluorescence.

#### unc-1(e1598)/unc9(e101)

The unc-9 mutation was verified with locomotion defective phenotype as well as PCR and restriction digest with BtsCI; the unc-1 mutation was verified by sequencing of the PCR product generated with primer pair KEB937/KEB938.

### Primer list

KEB836 CTCCTCATCCAGCGATCC KEB837 CCGCACGATTTGTGATGG KEB838 GTCTCTCTATCGGCCACC KEB839 CCAGATGCAAAGCCAGG KEB840 GTGTCGAGTGAAGGGAGC KEB911 CTTGGATGTATGATAATGCTC KEB912 GCACCTTTTCCAGCAATTAC KEB913 CTCTCTGATTGACATTCTTCC KEB914 CACCTATGTTAGAAACACGG KEB915 CCTCTCCGGAGATTACTTG KEB937 ACTGAACGGACTTTCTGCGA KEB938 GGGATCCAAATTTCAAAAGGTGC

## Supplementary Figure Legends

**S1:**
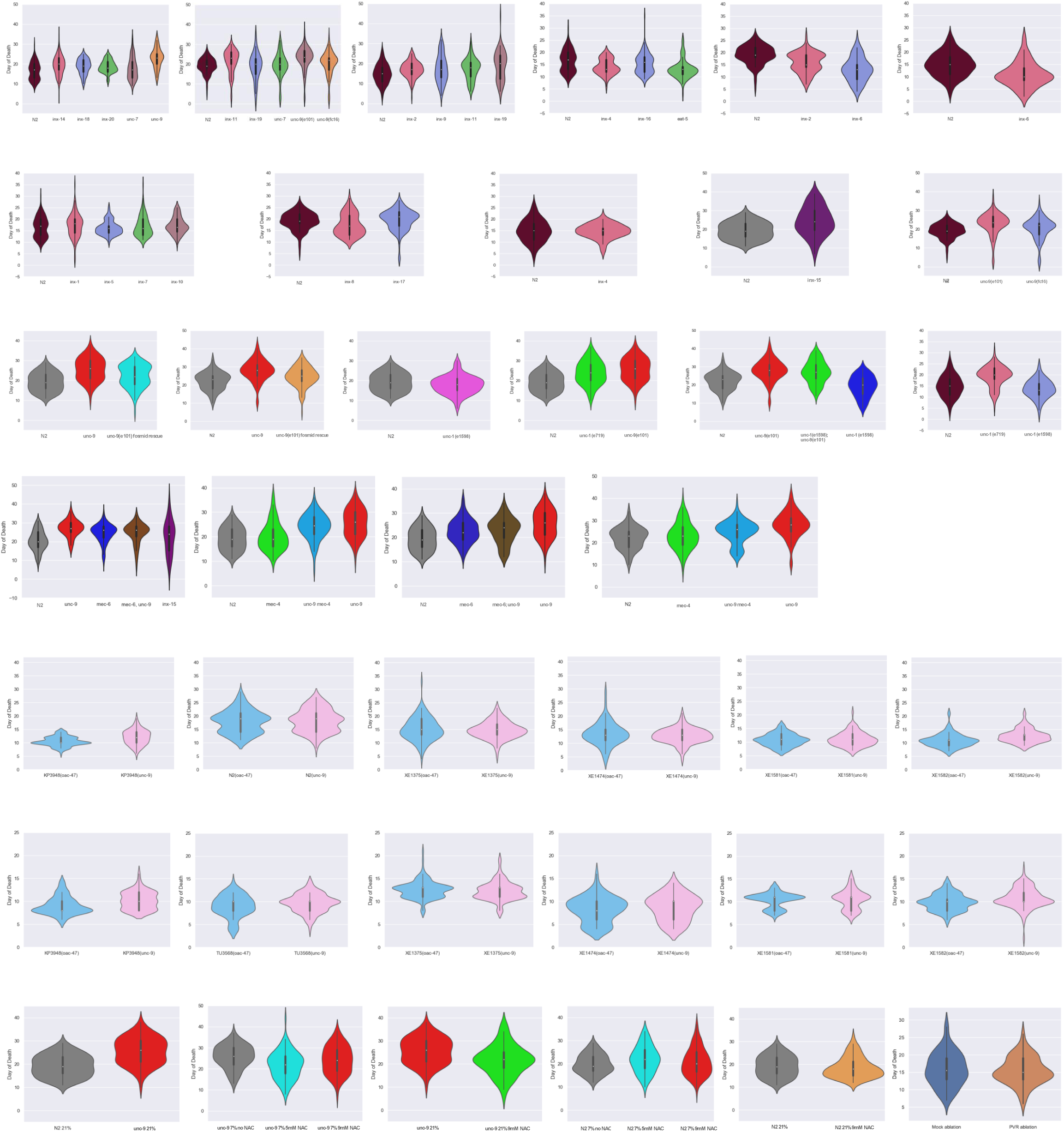
Violin plots representing the probability density of the lifespan data for all tested mutant and treatment conditions. The white dots represent the median, the thick black bars in the centre represent the interquartile range, and the thin black lines represent the rest of the distribution except for points that are determined to be outliers. Wider sections of the violin plot represent a higher probability that members of the population will take on the given value.

**S2:**
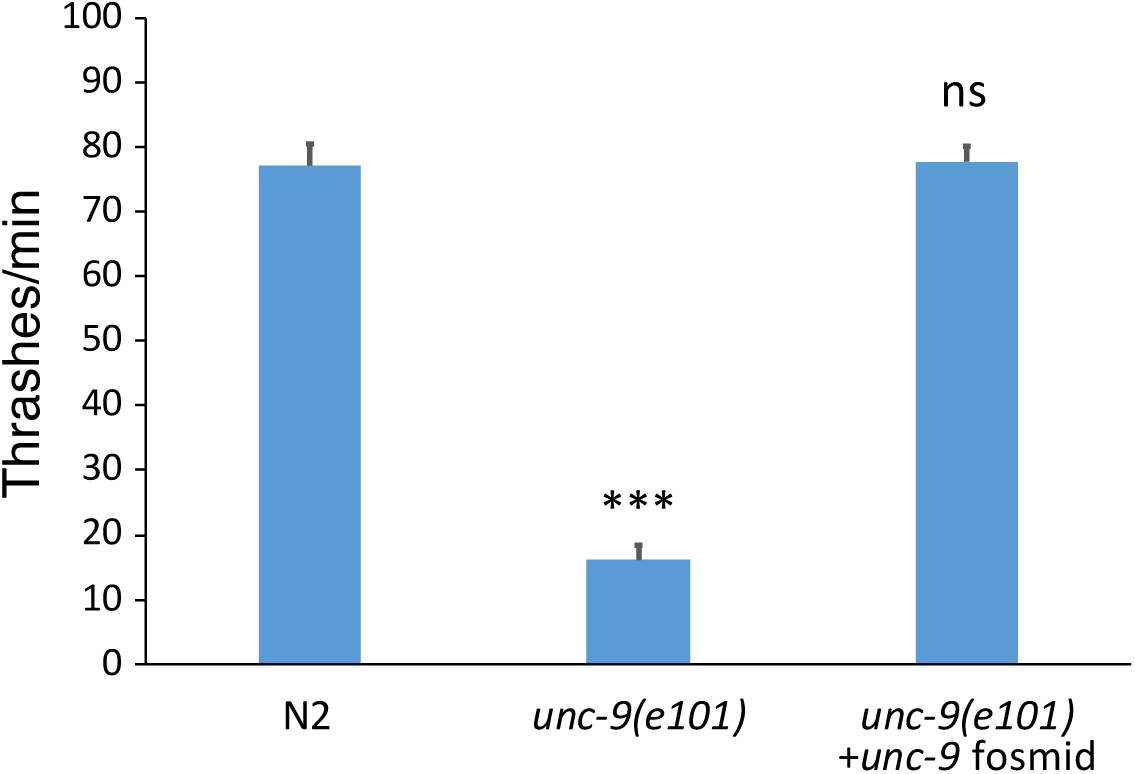
Lateral swimming movements (thrashes) per minute in N2 animals, *unc-9(e101)* mutants and *unc-9(e101)* mutants bearing a fosmid containing the *unc-9* genomic locus. ***P<0.001; ns not significant, using Student’s t-test.

**S3:**
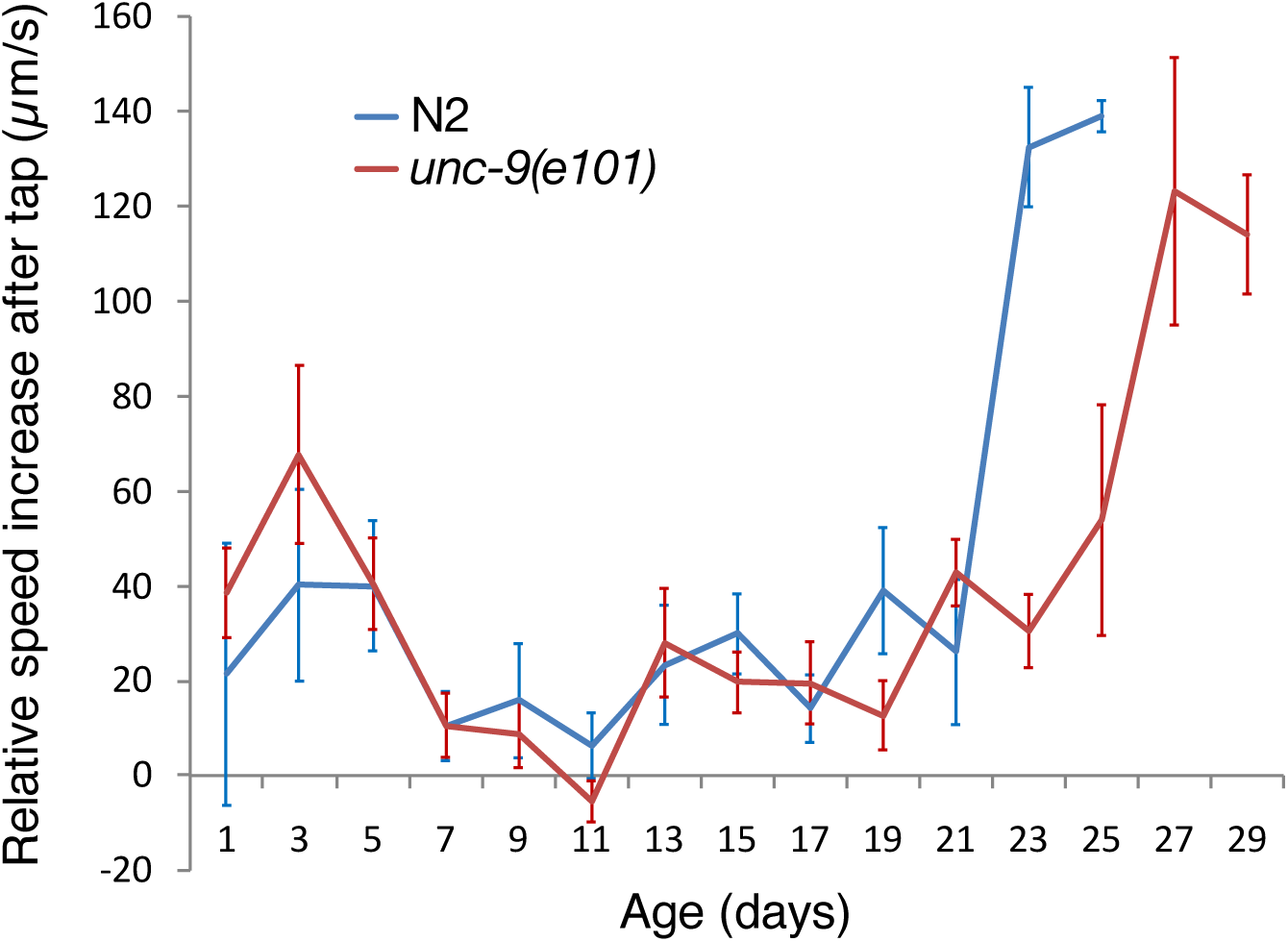
Changes in locomotory responses to a tap stimulus with age in *unc-9(e101)* and N2 animals. N=120 animals per group.

**S4:**
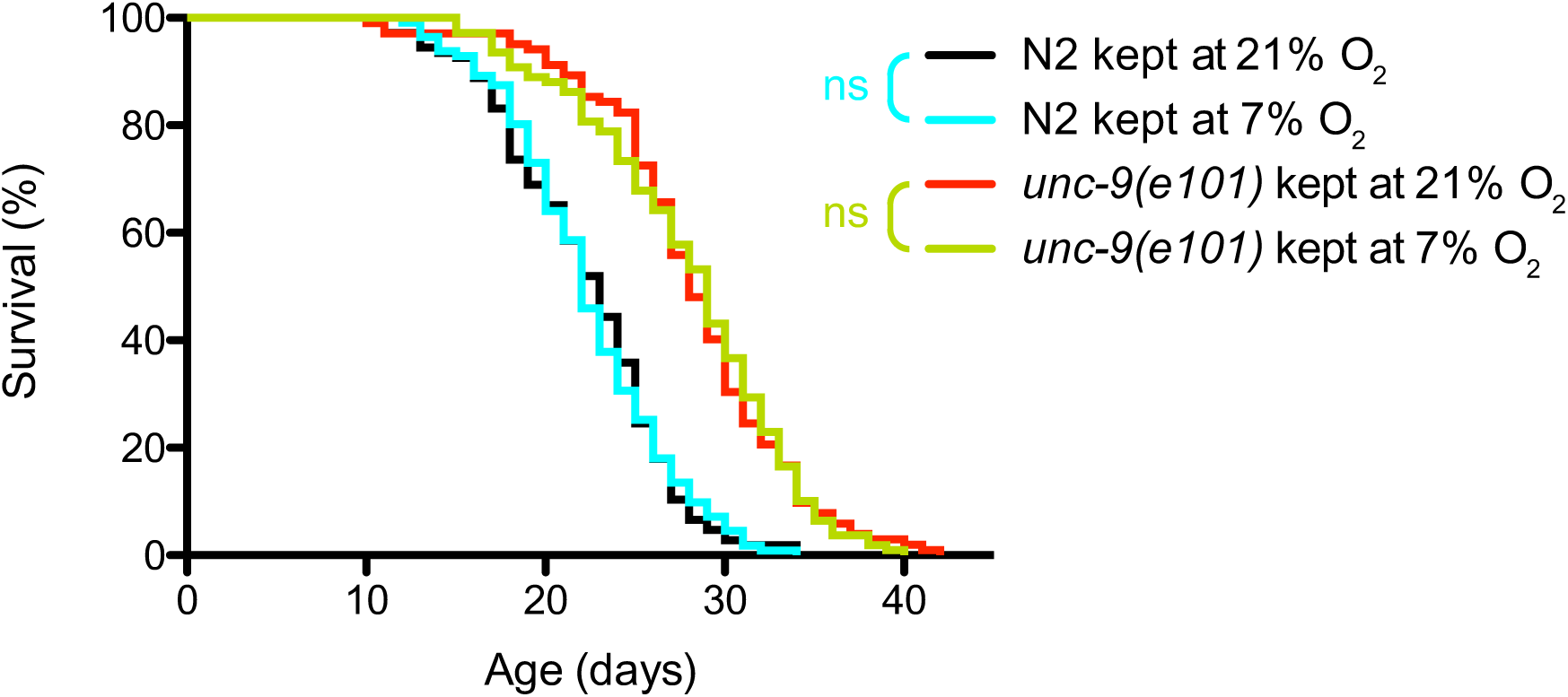
Lifespan of *unc-9(e101)* and N2 worms maintained at either 21% or 7% ambient O_2_. N=120 animals per group. ns not significant, using Log-rank test.

**S5:**
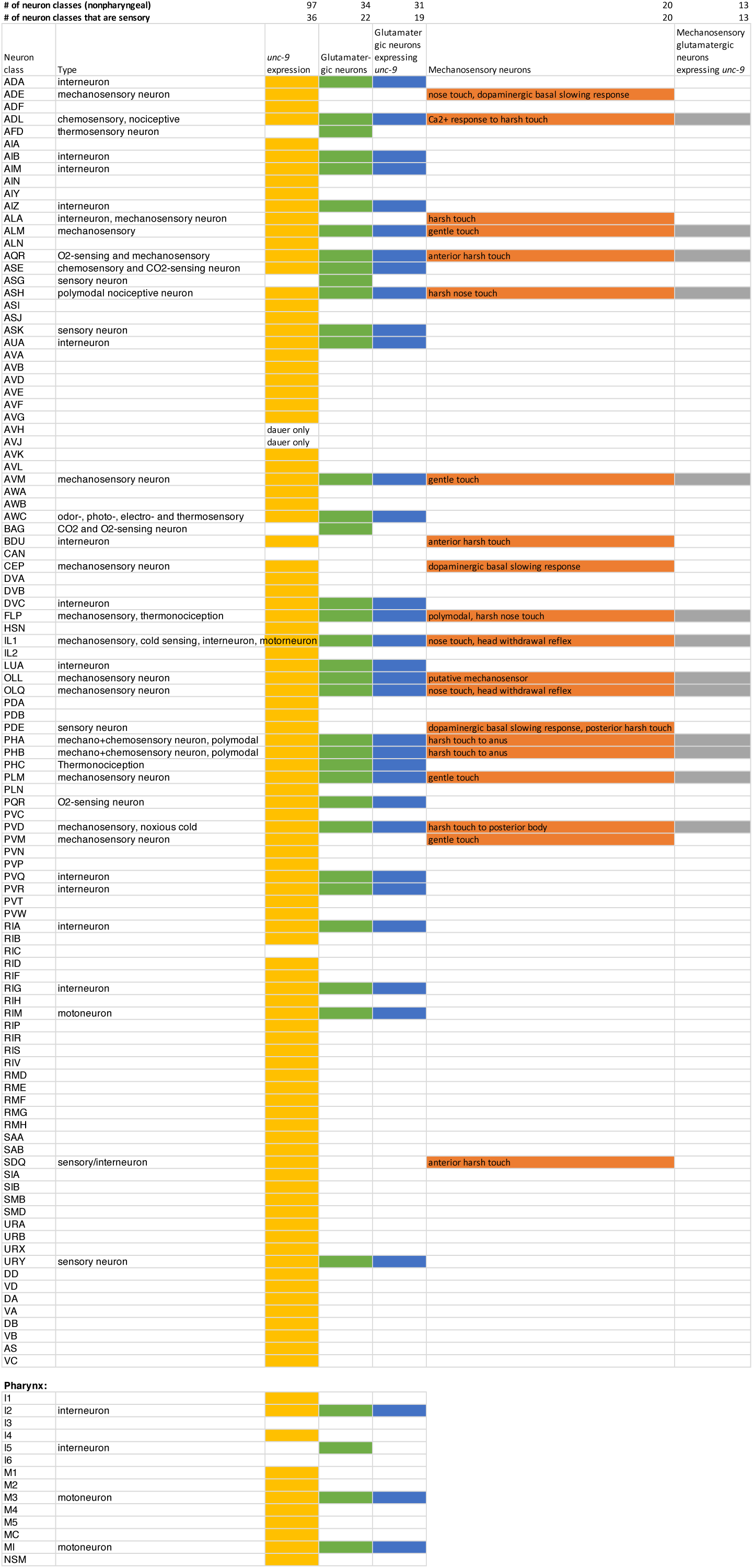
List of neuron classes in *C. elegans*, with neuron types indicated as classified in^1^. Classes with *unc-9* expression in non-dauer hermaphrodites are highlighted orange in the third column, based on^2^. The glutamatergic neurons (green) are based on *eat-4* expression^3^. Overlap of *unc-9* expression with glutamatergic and mechanosensory neurons is highlighted in blue and grey, respectively. The number of classes belonging to each category is given above the table. Information about the mechanosensory neurons is based on ^4–14^ and excludes stretch receptors.

**S6:**
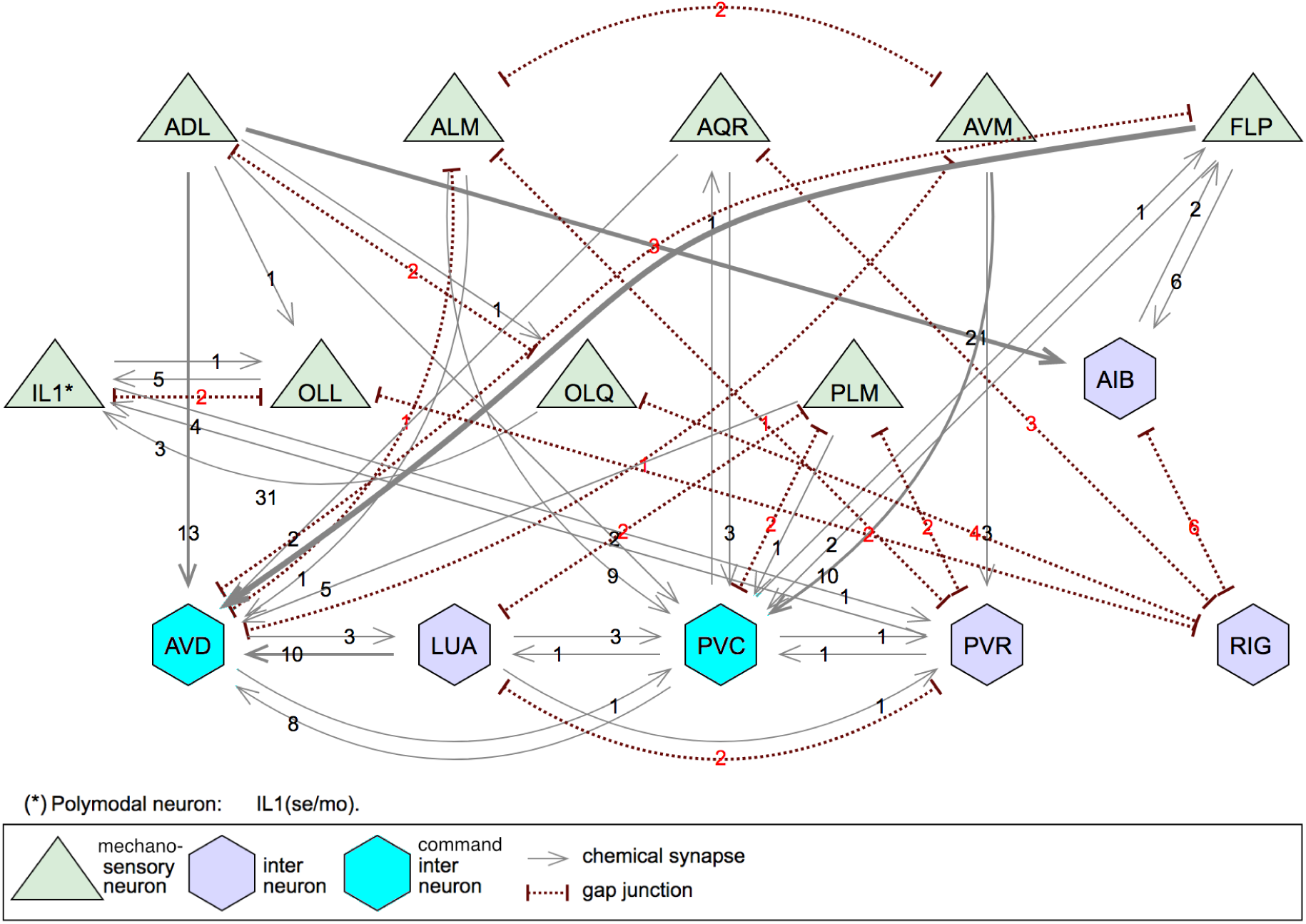
Chemical and electrical coupling of selected mechanosensory neurons, interneurons and command interneurons in *C. elegans,* based on the synaptic wiring described in^15^. The number of synapses between connected neurons is displayed.

**S7:**
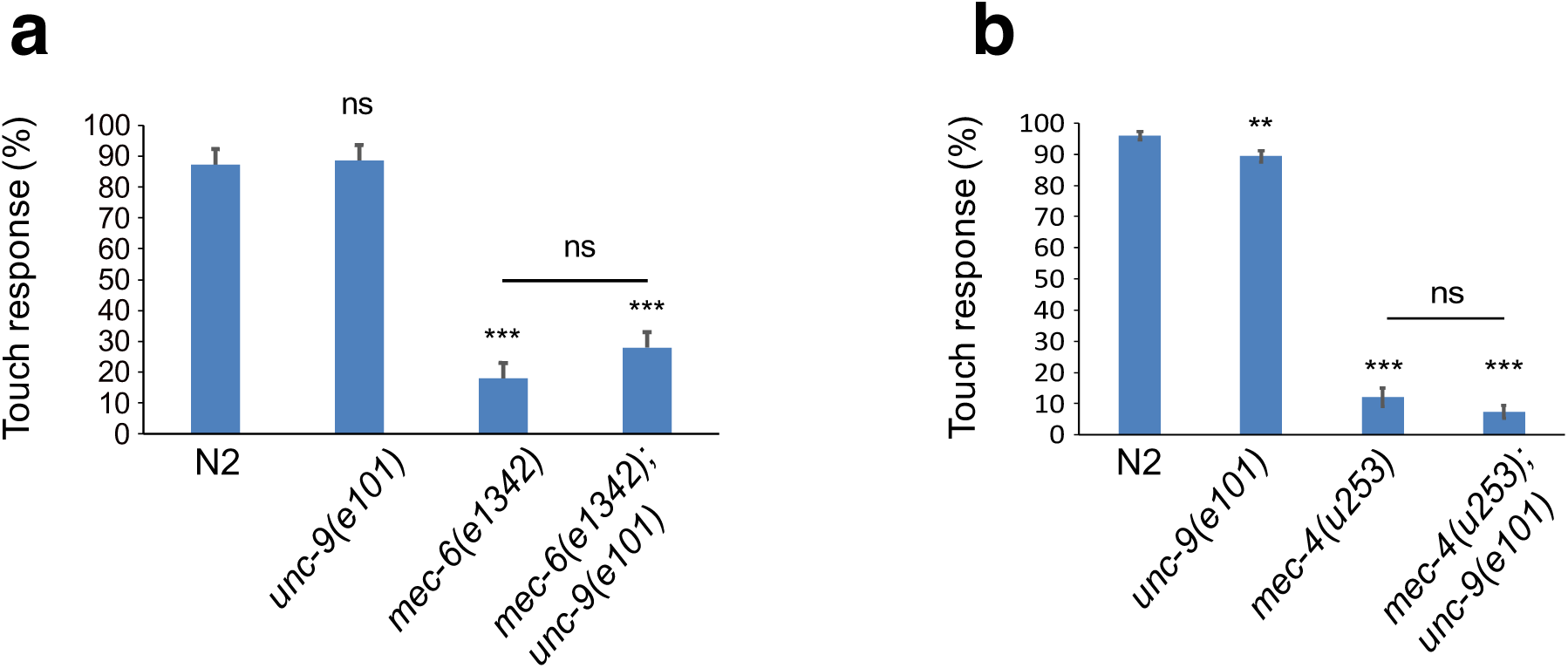
**a** Responses to touch with an eyelash pick in N2 and *unc-9(e101)* animals compared to *mec-6(e1342)* and *mec-6(e1342); unc-9(e101)*; and (**b**) *mec-4(u253)* and *mec-4(u253); unc-9(e101)* animals (N=15 animals per group). **P<0.01; ***p<0.001; ns not significant, using Student’s t-test.

**S8:**
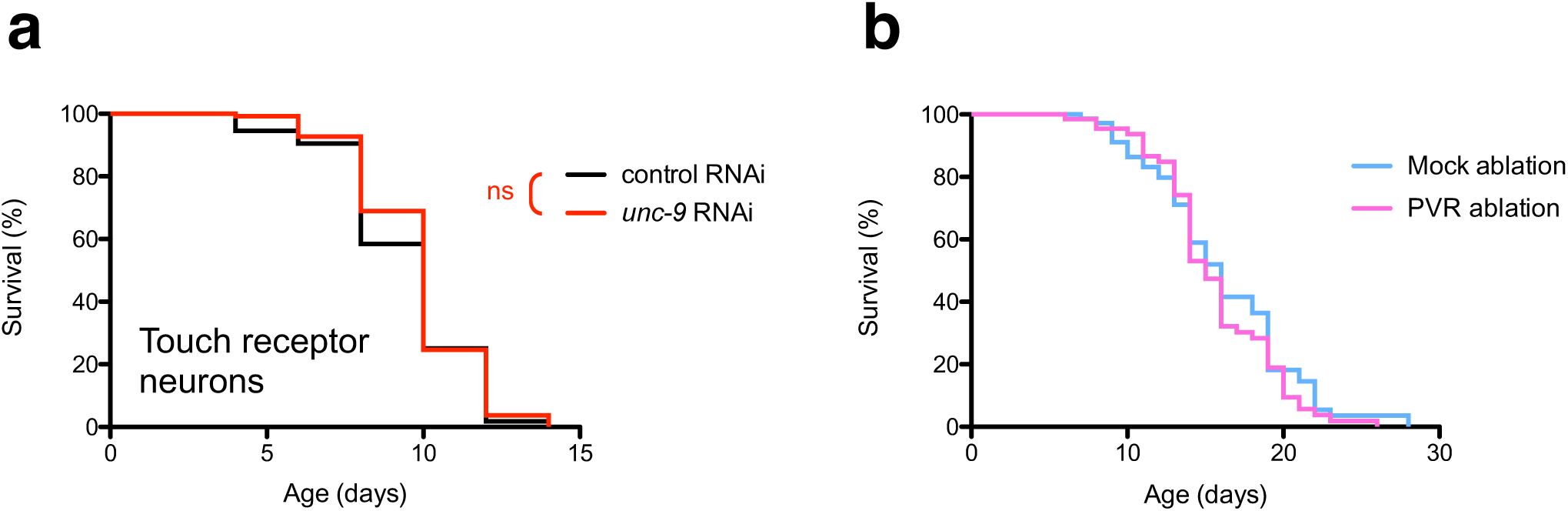
**a** Lifespan of controls and animals with *unc-9* knockdown by cell-specific RNA interference in touch receptor neurons; ns not significant, using Log-rank test. N=130 animals per group. **b** Lifespan of NY2054 animals where the PVR interneuron was laser ablated, versus controls that were mock ablated. ns not significant, using Log-rank test. N=79 animals per group.

**S9:**
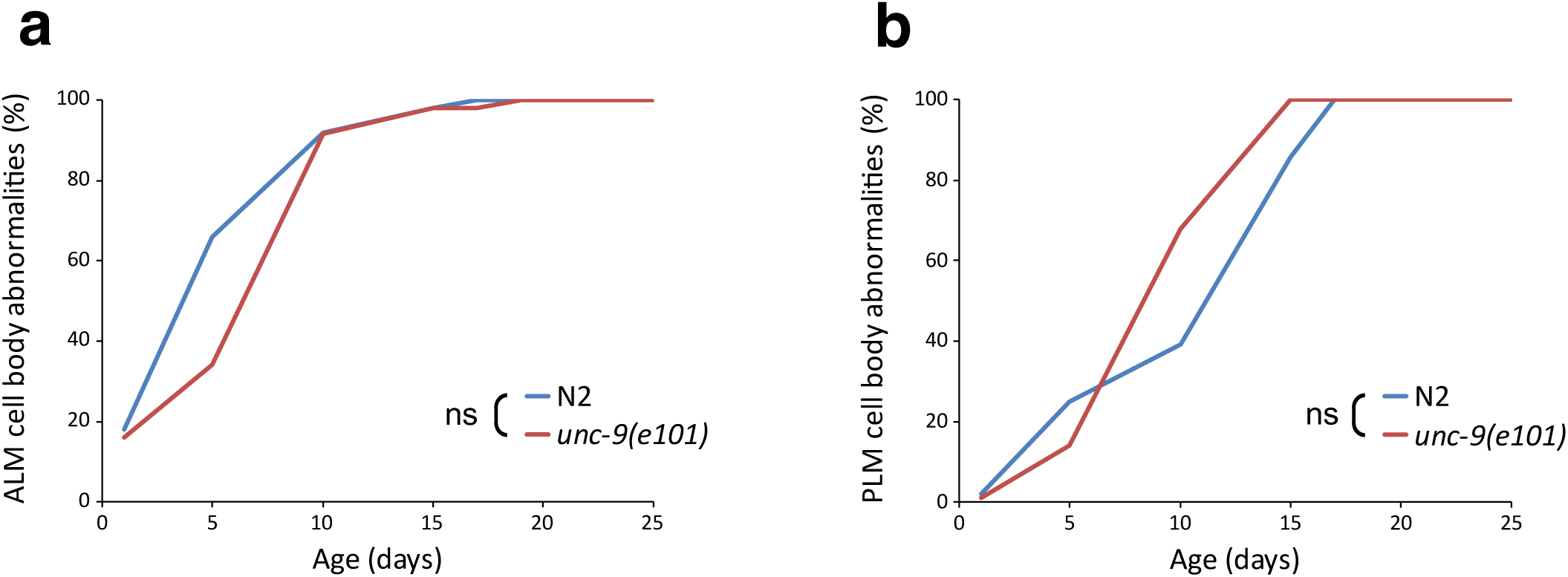
**a** Age-dependent decline of of ALM or (**b**) PLM cell body integrity is not significantly affected by genotype. N=50 animals per group. ns not significant, using a binomial regression model.

**S10:**
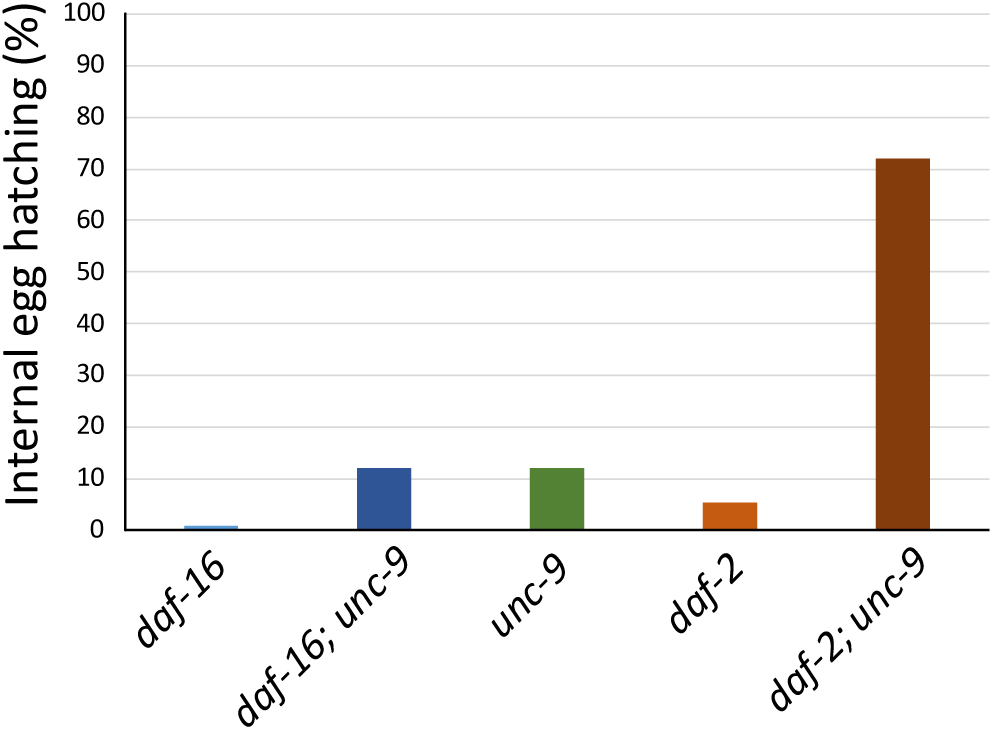
Propensity for internal egg hatching in *daf-16*, *daf-16; unc-9*, *unc-9*, *daf-2* and *daf-2; unc-9* mutants; N=120 animals per group.

**S11:**
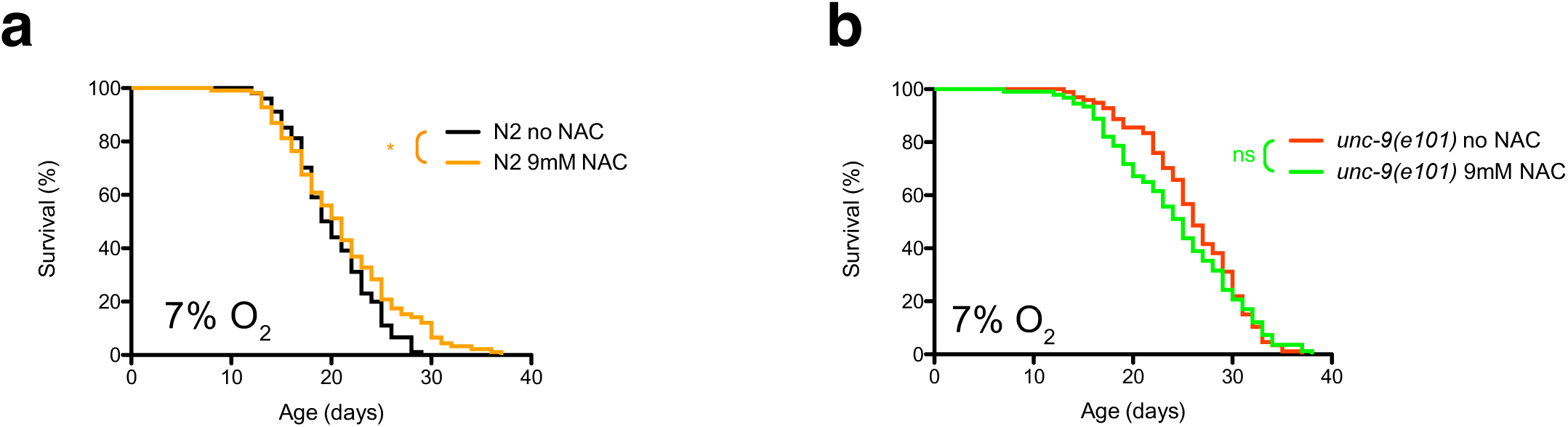
**a** Effect on lifespan of 9mM N-Acetyl-Cysteine (NAC) treatment versus untreated controls in N2 and (**b**) *unc-9(e101)* mutant animals, both maintained at 7% O_2_. N=120 animals per group. *P<0.05; ns not significant, using Log-rank test.

**Supplementary Table 1:** Information about the mutants used in the study

| Gene | Strain | Allele | Molecular lesion | Comment | Reference |
| --- | --- | --- | --- | --- | --- |
| *inx-1* | - | *tm3524* | 238 bp deletion | likely null, deletes exon 6 |  |
| *inx-2* | CX13325 | *ok376* | 992 bp deletion | almost certainly null, deletes most of exon 2 and all of exons 3 and 4 |  |
| *che-7* | RB1834 | *ok2373* | 1496 bp deletion | almost certainly null, deletes most of exon 3 and all of exons 4 and 5 |  |
| *inx-5* | RB1086 | *ok1053* | 1221bp deletion | putative null - 1221bp deletion in R09F10: 5058-6278; deletes all amino acids after pos. 285, takes out part of exon 5 and all of exon 6 | this study |
| *inx-6* | MR127 | *rr5* | P353L missense mutation | temperature sensitive allele, 25°C restrictive temperature | 18 |
| *inx-7* | RB1792 | *ok2319* | 1610 bp deletion | almost certainly null, deleted part of exon 1 and all of exons 2-5 in isoform a and exons 1-3 in isoform b |  |
| *inx-8* | VC116 | *gk42* | 850 bp deletion | likely loss-of-function or null allele, deletes exon 1; fertile. |  |
| *inx-9* | CX12726 | *ok1502* | 719 bp deletion | very likely null, deletes most of exons 2 and 3. Fertile; VC994 is 0x outcrossed |  |
| *inx-10* | RB2051 | *ok2714* | 689 bp deletion | likely null, most of exon 3 and part of exon 4 deleted |  |
| *inx-11* | RB2108 | *ok2783* | 752 bp deletion | likely null, exons 3, 4, 5 and part of 6 deleted (in isoform b) |  |
| *inx-14* | AU98 | *ag17* | R326H missense mutation | *ag17* is hypomorphic allele; *ok3267*, *tm2593* and *tm2864*, which are partial deletions, all marked as sterile. Arginine-to-histidine substitution in *inx-14* near the end of the predicted third transmembrane domain | 19 |
| *inx-15* | - | *tm3394* | 458bp deletion | likely null, most of exon 2 deleted |  |
| *inx-16* | EG144 | *ox144* | nonsense mutation at position 104 | likely null | 20 |
| *inx-17* | - | *tm3292* | 309 bp deletion | likely null - most of exon 2 and part of exon 3 deleted |  |
| *inx-18* | RB1896 | *ok2454* | 1704 bp deletion | putative null; 1704bp deletion, in C18H7: 30705-32408; deletes part of exon 4 and all of exon 5 | this study |
| *inx-19* | CX6161 | *ky634* | E54K/E70K missense mutation | described as loss-of-function allele | 21 |
| *inx-20* | CX12725 | *ok681* | 1483 bp deletion | likely null, deletes exons 2-8; see also RB683, bearing a different allele | 22 |
| *inx-20* | RB683 | *ok426* | 984 bp deletion | likely null, deletes part of exon 5 and 6-9 completely, part of exon 10 |  |
| *inx-21* | RB1929 | *ok2524* | 1960bp deletion | exons 3 and 4 deleted but is not a null allele (which would be sterile) - see <sup>23</sup> |  |
| *inx-22* | XM1011 | *tm1661* | 765bp deletion | likely null, deletes most of exon 2 and all of exon 3; viable and fertile | 23 |
| *eat-5* | DA1402 | *ad1402* | 1439 bp deletion | putative null deletion that removes exons 2 through 4. | 24 |
| *unc-9* | CB101 | *e101* | splice acceptor site mutation at last exon | reference allele, severe or null mutant | 25 |
| *unc-9* | CW129 | *fc16* | nonsense mutation at position 157 | putative null mutant resulting from premature stop in the intracellular loop between the second and third membrane-spanning domain | 25 |
| *unc-7* | CB5 | *e5* | nonsense mutation | likely null mutation, premature stop (Q216*) in the first predicted extracellular loop of UNC-7 | 26 |
| *unc-1* | CB719 | *e719* | frameshift at pos. 162 | putative null - a splice junction mutation causes frameshift and premature termination | 27 |
| *unc-1* | CB1598 | *e1598* | A216V missense mutation | canonical dominant gain-of-function allele | 28 |
| *mec-3* | CB1338 | *e1338* | A inserted at 2782 | frameshift, touch receptors do not differentiate, PVD nonfunctional | 29, 30 |
| *mec-4* | CB1611 | *e1611* | A713T substitution | dominant, all gentle-touch sensory neurons degenerate | 31 |
| *mec-4* | TU253 | *u253* | deletion in exon 3 | likely null, no mechanoreceptor currents | 32 |
| *mec-6* | CB1472 | *e1342* | W303* nonsense mutation | reference allele, nonsense mutation and therefore likely a null | 33 |

**Supplementary Table 3:** *C. elegans* strains used

| Strain | Genotype | Reference |
| --- | --- | --- |
| - | *inx-1(tm3524) X.* | 34 |
| CX13325 | *inx-2(ok376) X.* | 22 |
| VC260 | *inx-2(ok376) X.* | 35 |
| RB1834 | *che-7/inx-4(ok2373) V.* | 35 |
| RB1086 | *inx-5(ok1053) X.* | this study |
| MR127 | *inx-6(rr5) IV.* | 18 |
| RB1792 | *inx-7(ok2319) IV.* | 35 |
| VC116 | *inx-8(gk42) IV.* | 35 |
| CX12726 | *inx-9(ok1502) IV.* | 22 |
| VC994 | *inx-9(ok1502) IV.* | 35 |
| RB2051 | *inx-10(ok2714) V.* | 35 |
| RB2108 | *inx-11(ok2783) V.* | 35 |
| AU98 | *inx-14(ag17) I.* | 19 |
| - | *inx-15(tm3394) I.* | 34 |
| EG144 | *inx-16(ox144) I.* | 20 |
| - | *inx-17(tm3292) I.* | 34 |
| RB1896 | *inx-18(ok2454) IV.* | this study |
| CX6161 | *inx-19(ky634) I.* | 21 |
| CX12725 | *inx-20(ok681) I.* | 22 |
| RB683 | *inx-20(ok426) I.* | 35 |
| CX12725 | *inx-20(ok426) I.* | 22 |
| RB1929 | *inx-21(ok2524) I.* | 35 |
| XM1011 | *inx-22(tm1661) I.* | 23 |
| DA1402 | *eat-5(ad1402) I.* | 24 |
| CB101 | *unc-9(e101) X.* | 25 |
| CW129 | *unc-9(fc16) X.* | 25 |
| CB5 | *unc-7(e5) X.* | 26 |
| CB1338 | *mec-3(e1338) IV.* | 29,30 |
| CB1370 | *daf-2(e1370) III.* |  |
| CB1472 | *mec-6(e1342) l.* | 33 |
| CB1598 | *unc-1(e1598) X.* | 28 |
| CB1611 | *mec-4(e1611) X.* | 31 |
| CB719 | *unc-1(e719) X.* | 27,36 |
| CF1038 | *daf-16(mu86) l.* |  |
| KL126 | *unc-9(e101) X; cipEx29(WRM0611aH10 (Fosmid containing unc-9 genomic DNA); pmyo-2::mCherry)* | this study |
| KL177 | *daf-2(e1370) III.; unc-9(e101) X.* | this study |
| KL180 | *daf-16(mu86) l.; unc-9(e101) X.* | this study |
| KL197 | *Is[mec-3::GFP]; unc-9(e101) X.* | this study |
| KL230 | *unc-9(e101) X; cipEx29(WRM0611aH10 (Fosmid containing unc-9 genomic DNA); pmyo-2::mCherry)* | this study |
| KL240 | *mec-6(e1342) l; unc-9(e101) X.* | this study |
| KL243 | *mec-6(e1342) l; unc-9(e101) X.* | this study |
| KL246 | *cipEx29[WRM0611aH10(punc-9::unc-9; pmyo-2::mCherry)]* | this study |
| KL273 | *unc-9(e101) mec-4(u253) X.* | this study |
| KL276 | *unc-1(e1598) unc9(e101) X.* | this study |
| KP3948 | *eri-1(mg366) IV; lin-15B(n744) X.* | 37 |
| NY2054 | *ynIs54 [flp-20p::GFP] IV. ; him-5(e1490) V.* |  |
| TU253 | *mec-4(u253) X.* | 32 |
| - | *Is[pmec-3::GFP]* | 38 |
| TU3568 | *sid-1(pk3321) him-5(e1490) V; lin-15B(n744) X; uls71[(pCFJ90) myo-2p::mCherry + mec-18p::sid-1].* | 39 |
| XE1375 | *wpls36[unc-47p::mCherry] l; wpSi1[unc-47p::rde-1::SL2::sid-1 + Cbr-unc-119(+)] II; eri-1(mg366) IV; rde-1(ne219) V; lin-15B(n744) X.* | 40 |
| XE1474 | *wpSi6[dat-1p::rde-1::SL2::sid-1 + Cbr-unc-119(+)] II; eri-1(mg366) IV; rde-1(ne219) V; lin-15B(n744) X.* | 40 |
| XE1581 | *wpSi10[unc-17p::rde-1::SL2::sid-1 + Cbr-unc-119(+)] II; eri-1(mg366) IV; rde-1(ne219) V; lin-15B(n744) X.* | 40 |
| XE1582 | *wpSi11[eat-4p::rde-1::SL2::sid-1 + Cbr-unc-119(+)] II; eri-1(mg366) IV; rde-1(ne219) V; lin-15B(n744) X.* | 40 |
| ZM3087 | *unc-9(fc16) unc-7(e5) X.* | 41 |

